# Clearance of defective muscle stem cells by senolytics reduces the expression of senescence-associated secretory phenotype and restores myogenesis in myotonic dystrophy type 1

**DOI:** 10.1101/2022.06.22.497227

**Authors:** Talita C. Conte, Gilberto Duran-Bishop, Zakaria Orfi, Inès Mokhtari, Alyson Deprez, Marie-Pier Roussel, Isabelle Côté, Ornella Pellerito, Damien Maggiorani, Basma Benabdallah, Severine Leclerc, Lara Feulner, Jean Mathieu, Cynthia Gagnon, Gregor Andelfinger, Christian Beauséjour, Serge McGraw, Elise Duchesne, Nicolas A. Dumont

**Affiliations:** CHU Sainte-Justine Research Center. Montreal, QC, Canada; Department of pharmacology and physiology, Faculty of Medicine, Université de Montréal. Montreal, QC, Canada; Department of obstetrics and gynecology. Faculty of Medicine, Université de Montréal. Montreal, QC, Canada; Department of Health Sciences, Université du Québec à Chicoutimi, QC, Canada; Neuromuscular diseases interdisciplinary research group (GRIMN), Saguenay Lac-St-Jean integrated university health and social services hospital, QC, Canada; Department of Fondamental Sciences, Université du Québec à Chicoutimi, QC, Canada; CHU Sherbrooke Research Center, Faculty of Medicine and Health Sciences, Université de Sherbrooke, Québec, Canada; Department of Pediatrics, Faculty of Medicine, Université de Montréal. Montreal, QC, Canada; School of rehabilitation, Faculty of Medicine, Université de Montréal. Montreal, QC, Canada

## Abstract

Muscle weakness and atrophy are clinical hallmarks of myotonic dystrophy type 1 (DM1). Muscle stem cells, which contribute to skeletal muscle growth and repair, are also affected in this disease. However, the molecular mechanisms leading to this defective activity and the impact on the disease severity are still elusive. Here, we explored through an unbiased approach the molecular signature leading to myogenic cell defects in DM1. Single cell RNAseq data revealed the presence of a specific subset of DM1 myogenic cells expressing a senescence signature, characterized by the high expression of genes related to senescence-associated secretory phenotype (SASP). This profile was confirmed using different senescence markers *in vitro* and *in situ*. Accumulation of intranuclear RNA foci in senescent cells, suggest that RNA-mediated toxicity contribute to senescence induction. High expression of IL-6, a prominent SASP cytokine, in the serum of DM1 patients was identified as a biomarker correlating with muscle weakness and functional capacity limitations. Drug screening revealed that the BCL-XL inhibitor (A1155463), a senolytic drug, can specifically target senescent DM1 myoblasts to induce their apoptosis and reduce their SASP. Removal of senescent cells re-established the myogenic function of the non-senescent DM1 myoblasts, which displayed improved proliferation and differentiation capacity *in vitro*; and enhanced engraftment following transplantation *in vivo*. Altogether this study presents a well-defined senescent molecular signature in DM1 untangling part of the pathological mechanisms observed in the disease; additionally, we demonstrate the therapeutic potential of targeting these defective cells with senolytics to restore myogenesis.

## INTRODUCTION

Myotonic dystrophy type 1 (DM1) is a rare disease that affects approximately 1:8,000 individuals in the world^1, 2^. DM1 is caused by an unstable (CTG) nucleotide repeat in the *dystrophia myotonica protein kinase* (*DMPK*) gene^3^. CTG repeats in DNA lead to toxic mRNA accumulation causing splicing defects mainly attributable to binding protein sequestration. In healthy individuals, the *DMPK* alleles have 5 to 37 CTG repeats, but in DM1, there can be hundreds to thousands of these repeats; with the number of repeats partially correlating with the age of onset and the severity of the disease^4–7^. Patients are classified into five clinical phenotypes according to the CTG expansion size, and the occurrence and onset of their main symptoms: congenital, infantile, juvenile, adult and late-onset^8^.

DM1 is a multisystemic disease that affects different organs, and particularly the skeletal muscles, leading to progressive weakness and atrophy^9^. Muscle stem cells (MuSC), which contribute to skeletal muscle growth and repair are also affected in this disease^5^. In healthy condition, quiescent MuSC are activated after an injury and become myoblasts that proliferate extensively before exiting cell cycle to self-renew or to differentiate and fuse to form new myofibers^10^. Previous studies performed on mixed cell populations (non-purified by FACS) extracted from skeletal muscle samples of DM1 patients showed that these cells exhibit signs of cellular senescence (a state of irreversible cell cycle arrest), such as enzymatic senescence-associated β-Galactosidase (SA-β-Gal) activity, expression of the cell cycle inhibitor p16, loss of nuclear integrity, and nuclear envelope invagination^11, 12^. It was shown using *in vitro* models of DM1 that cellular senescence is independent from telomere shortening but is rather induced by the activity of the cell cycle inhibitor p16 and/or p21 in response to CTG-related stress^12–14^. This increase in cellular senescence is associated with reduced proliferative capacity of DM1 myoblasts and delayed differentiation and fusion into myofibers^5, 12, 14^. Overexpression of cyclin-dependent kinase 4 (Cdk4) to bypass p16-induced senescence restored the proliferation of DM1 cells to the level of healthy control cells^12^.

In addition to decreasing the pool of competent cells, senescent cells secrete a broad range of molecules including proinflammatory cytokines (e.g., IL-1, IL-6) and matrix proteinases (e.g., MMP1, MMP3). This cocktail, known as the Senescence-Associated Secretory Phenotype (SASP), alters the microenvironment and leads to detrimental changes in neighbouring cells and in the whole organism^15, 16^. Particularly, IL-6 is associated with muscle wasting conditions and promote the exhaustion of the MuSC pool^17–19^. Chronic exposure to SASP is considered as one of the major sources of deleterious effects on tissue maintenance and regenerative capacity^15, 20^; however, its impact on DM1 pathogenesis has not been investigated yet.

Considering that senescent cells accumulate with time and have detrimental impacts on neighbouring cells, they represent an attractive target for the treatment of many disorders^21^. Pioneer studies showed that genetic removal of senescent cells expressing the cell cycle inhibitor p16 in aging mice enhances their lifespan^20, 22^. Moreover, pharmacological compounds, called senolytics, can selectively eliminate senescent cells^23^, which reduces the production of frailty-related proinflammatory cytokines in aged mice, and improves physical function and lifespan^24^. Considering that DM1 has been described as a progeroid syndrome (i.e. mimicking premature aging)^25^, and that skeletal muscles in DM1 share many similarities with aged muscles such as cellular senescence, MuSC defects, and muscle wasting^26^, it suggests that senolytics hold great therapeutic potential for the treatment of DM1. However, the impact of senolytics in muscular diseases such as DM1 has never been investigated.

Here, we generated the first single-cell transcriptomic atlas of DM1 to identify the molecular signature of myogenic cell subpopulations. Using myoblasts isolated from DM1 patients and age- and sex-matched healthy controls, we identified a specific subset of senescent-like cells expressing high levels of SASP. We confirmed cellular senescence using different markers (p16, p21, SA-β-Gal) in primary myoblasts of DM1 patients *in vitro* and in skeletal muscle biopsies *in situ*. Furthermore, we demonstrated that the level of IL-6, a key SASP cytokine, is correlated with upper- and lower-limb muscle weakness as well as functional capacity limitations in patients affected by DM1. Thereafter, we screened different senolytic drugs and have identified the B-cell lymphoma-extra large (BCL-XL) inhibitor, A1155463, as the most promising molecule to eliminate specifically senescent myoblasts cultured from DM1 patients’ samples, without affecting cells cultured from healthy controls’ samples. Elimination of senescent cells reduced SASP expression and restored the myogenesis potential of the myoblasts *in vitro* and following cell transplantation *in vivo*. Altogether, these findings support the importance of cellular senescence in the disease severity, and the therapeutic potential of senolytics to target defective myogenic cells and reduce SASP expression in DM1.

## RESULTS

### Single-Cell RNAseq identifies a subset of senescent cells in DM1 myoblasts

Considering that cellular senescence only affects a subset of cells at a given moment, we took advantage of the single-cell RNA sequencing (scRNAseq) technology to identify the distinct transcriptional profile in cellular subpopulations of DM1 myoblasts. Myoblasts were sorted by FACS (Suppl. Fig. 1) and purity was confirmed by immunofluorescence (Suppl. Fig. 2). Single-cell transcriptomic sequencing was performed on pooled myoblasts at the same early passage (P4) from 3 DM1 patients (juvenile phenotype) and 3 age- and sex-matched healthy controls (Suppl. Table 1). After filtering and removal of doublets in the sequencing data, we analysed 1,613 control cells and 1,714 DM1 patient cells (Suppl. Fig. 3). Both controls and DM1 cells express high levels of myogenic cell markers such as *CD82, ITGB1 (CD29), MYF5*, *MYOD1*, *NCAM1* (*CD56*), and *DES*, as well as low levels of the fibroadipogenic (*PDGFRa* and *CD34*), endothelial (*CD31* and *TIE1*), and myeloid (*CD11B* and *CD45*) cell markers (Suppl. Fig. 4). UMAP plot of merged control and patient cells revealed that these cells cluster differently (Fig. 1A). Differential expression analysis revealed that the expression of 516 genes was significantly changed (P < 10e-6; n=266 downregulated and n=250 upregulated) in DM1 patients compared to controls (Fig. 1B, Suppl. Table 2). Compared to healthy control cells, DM1 cells showed lower expression in genes associated to myogenic markers (e.g., *MYF5, MYOD1, MYF6, SIX1*), extracellular matrix components (e.g., *COL1A1, COL1A2, COL3A1, COL5A2, LAMA2, FN1*), Notch signalling (e.g., *HEYL, RBPJ*), and cell cycle (e.g., C*ENPF, CCNA2, CCNB1, CCNB2, CDK1, MKI67*) (Fig 1B,C; suppl. table 2). Downregulated genes in DM1 cells were enriched for biological processes related to cell cycle, muscle structure development, and extracellular matrix organization (Fig 1D,F). Top upregulated genes in DM1 cells included cytokines/chemokines related to inflammation (e.g., *CXCL1, CXCL2, CXCL3, CXCL5, CXCL6, CXCL8, IL1B, IL6, CSF3*), and metalloproteinases (e.g., *MMP1, MMP3*) (Fig 1 B,C; suppl. table 2). Upregulated genes in DM1 cells were enriched for biological functions included cytokine signaling, cellular response to stress, autophagy, apoptosis, and senescence (Fig 1E,G). SCENIC analysis was used to predict transcription factors regulating the gene sets. Results showed that downregulated genes are controlled by different transcriptions factors related to myogenesis (e.g., MYF5, MYOD1, MYF6, SIX1), while for the genes upregulated in DM1 the transcriptions factors related to the NF-kB pathway (e.g., NFKB1, NFKB2, REL, RELB) were identified (Fig. 1H,I).

**Figure 1:**
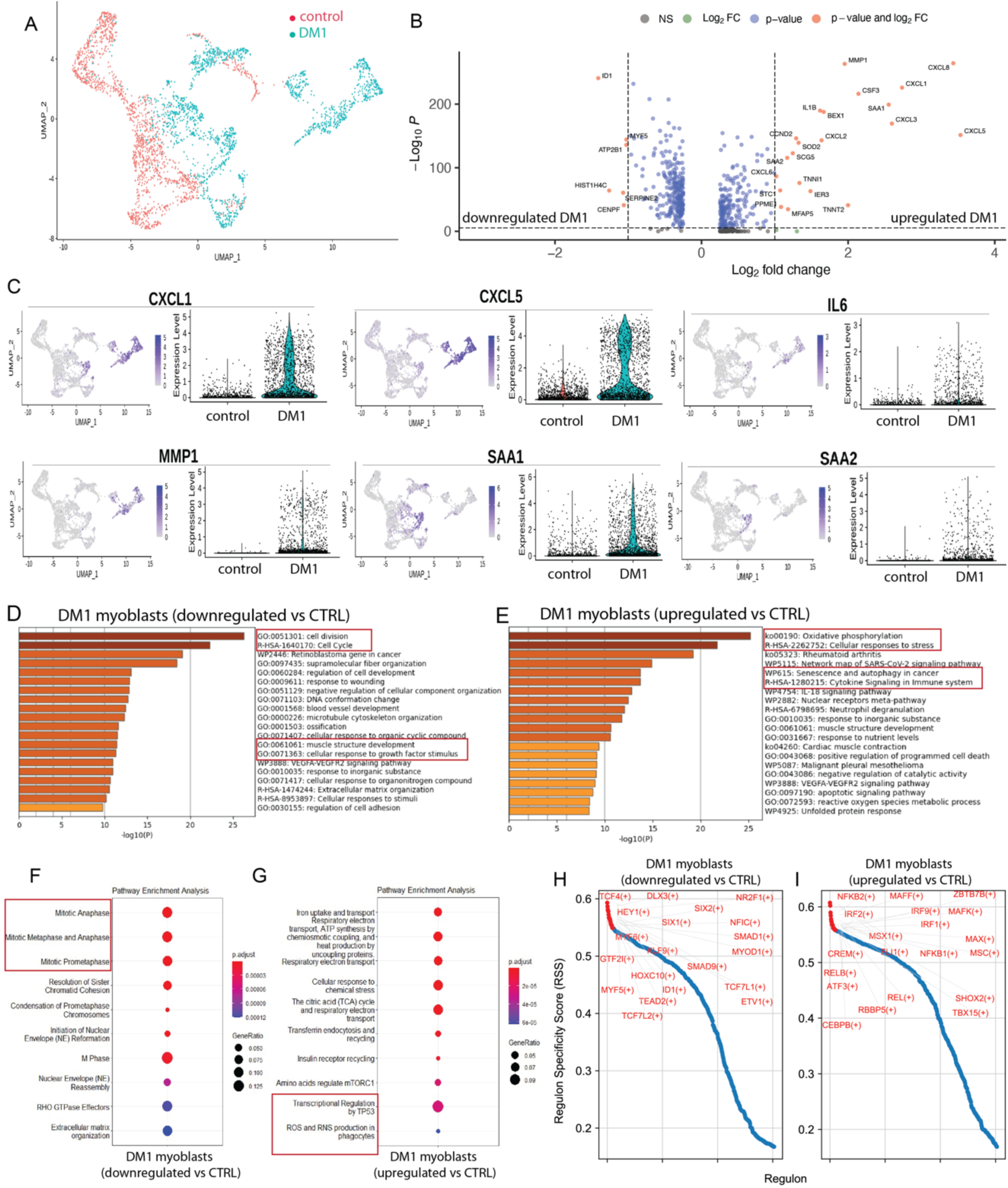
scRNA-seq analysis of myogenic cells from control and DM1 patients. **A)** UMAP embeddings displaying the control (red dots) and DM1 patient cell populations (blue dots). **B)** Volcano plot depicting differentially expressed genes (DEGs) downregulated or upregulated in DM1 patient cells vs controls. The x-axis represents the log2 of the expression fold change and the y-axis represents the negative log10 of the adjusted p -value of the Wilcoxon rank-sum test. Blue dots represent the transcripts that were statistically significant (P < 10e-6; dotted horizontal line) while red dots represent the top DEGs (|log2FC| > 1.0; vertical dotted line). **C)** Feature and violin plots showing the expression of selected DEGs in DM1 patient and control cells. **D,E)** Functional enrichment analysis with GO Biological Processes analysis using DEGs downregulated (**D**) and upregulated (**E**) in DM1 vs control (CTRL) cells. The X-axis represents the –log10(P-value). **F,G)** Bubble diagram showing the top 10 Reactome pathways for DEGs downregulated (**F**) and upregulated (**G**) in DM1 vs control cells. Vertical axes show enriched terms, and horizontal axes represent the genes in each cluster. Larger node size represents the larger ratio of enriched genes/total genes. **H,I**) Rank for regulons for downregulated genes (**H**) or upregulated genes (**I**) in DM1 vs control cells based on regulon specificity score (RSS).

UMAP plot showed that cells expressing high levels of pro-inflammatory cytokines (e.g., *CXCL1, CXCL8, IL6*) in DM1 patients are restricted to specific cell clusters (Fig. 1C). Further, *SAA1* and *SAA2*, two target genes of pro-inflammatory molecules such as IL-1, IL-6, and TNFα^27^, are also upregulated in specific subsets of DM1 myogenic cells. To further explore this senescent-like phenotype we compared the different clusters to gene sets related to cellular senescence and SASP production^28^. Based on this profile, we were able to classify the DM1 cells in 3 subpopulations: one subset of fully senescent cells expressing high levels of senescent genes and SASP (clusters 2, 3, 6), one subset of early-senescent cells expressing senescence markers that did not transcribe into SASP cytokine secretion (cluster 4), and one subset non-senescent cells (clusters 0, 1, 5) (Fig. 2 A-D). By comparing the molecular signature of the senescent and non-senescent clusters versus all the combined clusters found in our control sample, we observed a clear senescent molecular signature present only in the senescent cluster (*CXCL1, CXCL3, CXCL5, CXCL8, MMP1, SOD2*); and that the non-senescent cluster in DM1 samples expressed a similar molecular profile as our control sample (Fig. 2E).

**Figure 2:**
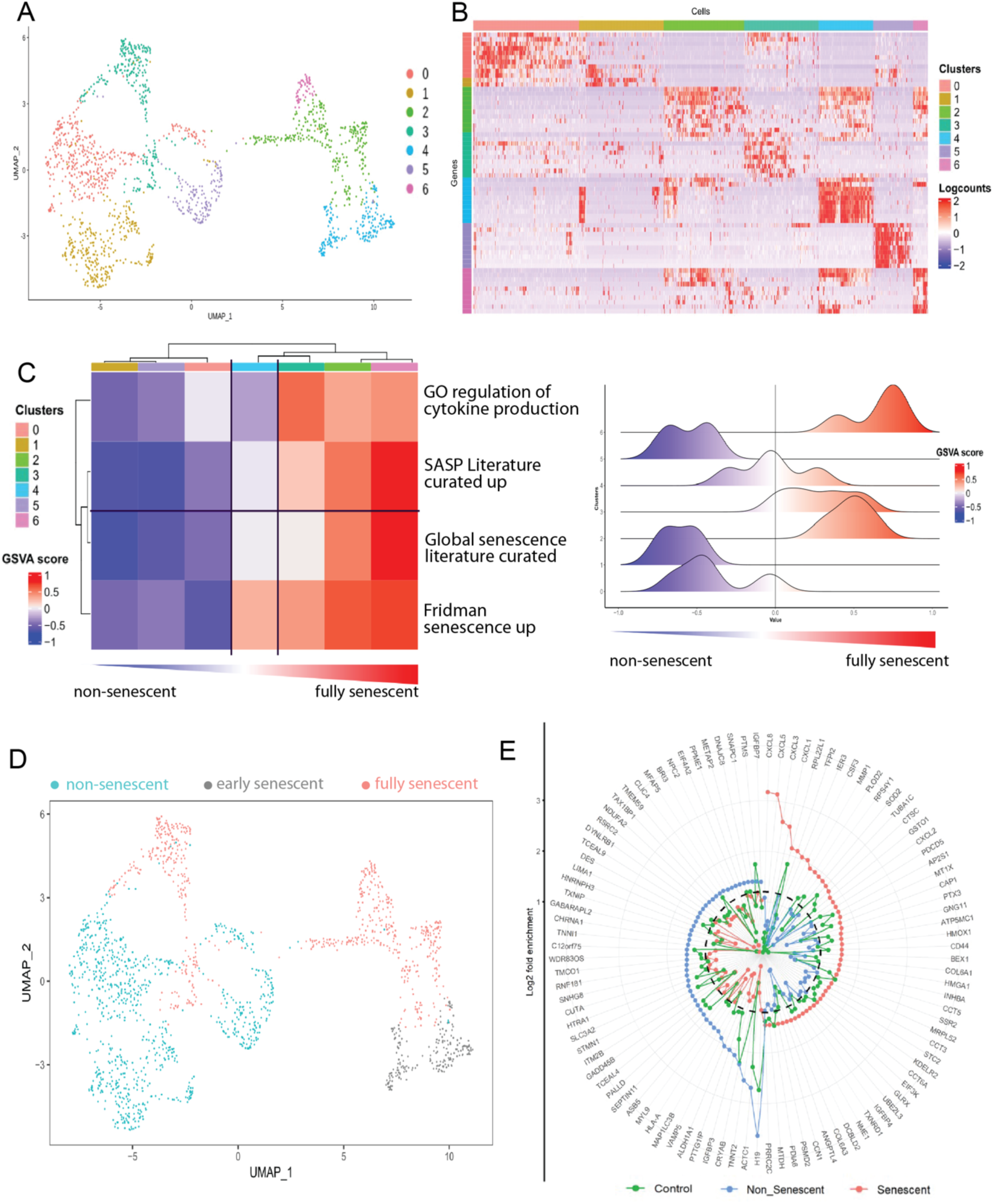
Identification of senescent cell subpopulations in patients with DM1. **A)** UMAP visualization showing unsupervised clustering revealing 7 distinct cell populations in patient cells. **B)** Heatmap showing the top differentially expressed genes (DEGs) from each cluster. **C)** Heatmap and ridge plots showing the fold change of GSVA score on gene sets related to cellular senescence and cytokine secretion comparing DEGs transcripts from each cluster. Clusters with positive GSVA values for all the gene sets were annotated as fully senescent while clusters with mixed and negative GSVA scores were annotated as early-senescent and non-senescent respectively. **D)** UMAP embedding characterizing the fully senescent, early-senescent, and non-senescent populations in patient cells. **E)** Radar plot of DEGs of senescent (red) versus non-senescent (blue) cells. Expression of the DEGs in the control cells are shown in green. Y-axis represents the log2 of the expression fold change. Dotted line delineates the threshold of log2FC = 1.2.

Production of SASP by senescent cells varies depending on the cell type and inducer^29, 30^. Thus, we selected all the top differentially expressed genes (>1.7-fold) in DM1 and controls samples and we investigated the presence of these genes in the SASP Atlas^30^, a proteomic database that analysed the SASP factors secreted by different cell types (fibroblasts and epithelial cells) subjected to multiple senescence inducers (irradiation, oncogene-induced, treatment-induced). These findings indicate that the top genes overexpressed in DM1 patients (e.g., *STC1, SOD2, MMP1, CXCL5, CXCL8*) are canonical SASP markers expressed at the protein level by senescent cells, regardless of the cell type or the senescence inducer (Suppl. Fig. 5A). On the other hand, the top upregulated genes in healthy control myoblasts (e.g., *COL3A1, TOP2A*) are negative regulators of cellular senescence in the SASP Atlas (Suppl. Fig. 5B).

### Cellular senescence is associated with reduced myogenesis in DM1

As single-cell analysis revealed distinct molecular signatures associated to cellular senescence in DM1 cells, we further explore how this process was affected in myogenic cells *in vitro* and *in situ*. First, we performed SA-β-Gal assay on myoblasts *in vitro*, and we observed an 11-fold increase (p = 0.0023) in the proportion of β-Gal + senescent cells in patient-derived DM1 myoblasts compared to healthy controls (Fig. 3A-B). Further expression analysis on cultured myoblasts confirmed that *P16* is significantly upregulated (p = 0.046) in DM1 compared to controls (Fig. 3C). As senescence is characterized by cell cycle arrest, myoblast proliferation was assessed using a live cell imaging system (IncuCyte), and the results showed a significant reduction (p = 0.042) in cell proliferation in DM1 patients compared to controls (Fig. 3D).

**Figure 3:**
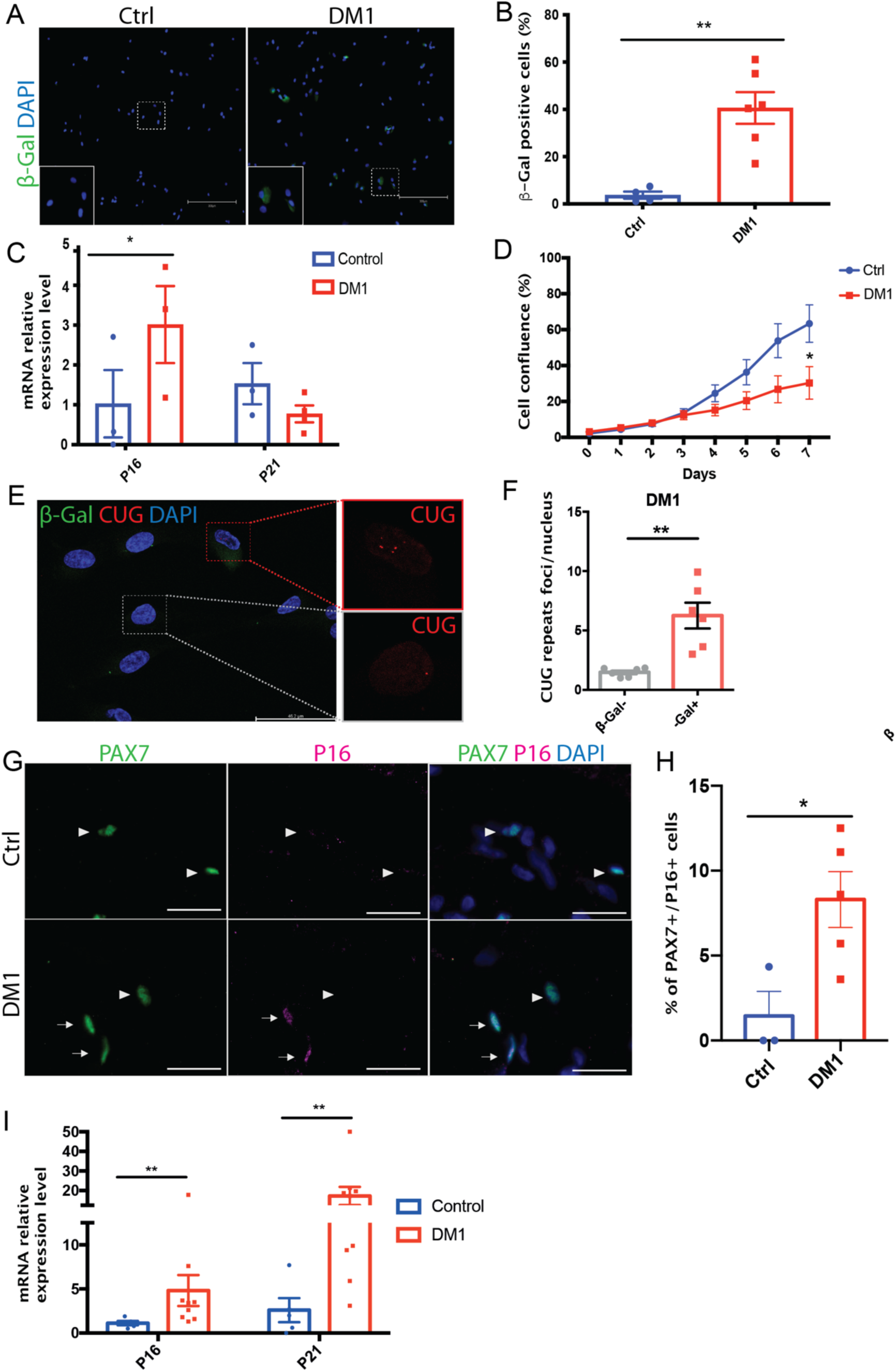
MuSC senescence is a hallmark of DM1 *in vitro* and *in situ*. **A)** Representative micrographs of primary myoblast culture of one control individual (right image) and one DM1 patient (left image) enzymatically stained for AlexaFluor 488 SA-ß-Gal to identify senescent cells (green). Scale bar: 300 μm. **B)** Quantification of the percentage of SA-ß-Gal+ cells (n = 4-6); *\*\*p = 0.0023*. **C)** Quantitative real-time PCR for the senescence markers *P16* and *P21* in primary myoblasts from controls and DM1 patients (n = 3-4); *\*p = 0.046.* **D)** Growth curve of control and DM1 myoblasts cultured *in vitro* for 7 days; *\*p = 0.042.* **E)** Representative micrograph of co-immunostaining for RNA FISH (CUG repeats, red), SA-β-Gal (green), and DAPI (blue). Red dashed lines identify a senescent cell and grey dashed lines a non-senescent cell. **F**) Quantification of the number of intranuclear RNA foci per senescent or non-senescent cell in DM1 myoblasts. *\*\*p = 0.0017.* **G**) Co-immunofluorescence labeling of PAX7 (green, MuSC marker) and P16 (magenta, senescence marker) on muscle sections of a control (upper images) and a DM1 patient (bottom images). White arrowheads indicate PAX7+P16-MuSC and white arrows indicate PAX7+P16+ senescent MuSC. Scale bars: 50 μm. **H)** Quantification of senescent MuSC expressing PAX7 and P16; *\*p = 0.0219.* **I)** Quantitative real-time PCR for the senescence markers P16 (*\*\*p = 0.0075*), and P21 (*\*\*p = 0.004*) on muscle biopsies from controls and DM1 patients (n = 5-9). Data are expressed as means +/-SEM.

To determine if the accumulation of toxic RNA is the cause of cellular senescence in DM1, the number of intranuclear RNA foci was compared in senescent (SA-β-Gal+) and non-senescent (SA-β-Gal-) DM1 myoblasts. A 4-fold increase (p = 0.0017) was observed in the number of RNA foci/cell in senescent DM1 cells compared to non-senescent cells (Fig. 3E,F, and suppl. Fig. 6).

Thereafter, cellular senescence markers were assessed on flash frozen skeletal muscle biopsies to determine if senescence is also a feature of the disease *in situ*. Immunofluorescence staining on skeletal muscle section showed a 5-fold increase (p=0.022) in the proportion of satellite cells (PAX7+) co-expressing the cell cycle inhibitor P16 (Fig. 3G,H) in DM1 muscles compared to controls. Gene expression analysis of skeletal muscle biopsies confirmed a significant upregulation in the expression of the cell cycle inhibitors *P16* (5-fold; p=0.0075) and *P21* (17-fold; p=0.004) in samples from DM1 patients compared to controls (Fig. 3I).

### SASP production is correlated with muscle weakness in DM1 patients

To determine the impact of SASP production on muscle function *in vivo*, blood serum from DM1 patients (n=103; suppl. table 3) was collected to measure the level of expression of IL-6, a prominent SASP cytokine. Thereafter, the maximum isometric muscle strength of different muscle groups of the lower limb (ankle dorsiflexors, hip flexors, knee extensors, knee flexors) and upper limb (shoulder flexors, shoulder abductors, elbow flexors, elbow extensors) was quantitatively assessed. Using equations that adjust for age, sex, and weight, the predicted strength value of each muscle group was calculated, which allowed to identify a relative deficit compared to normative value^31^. For the lower limb, a significant negative correlation between IL-6 levels and predicted maximal muscle strength was observed for ankle dorsiflexors (R: −0.426; p<0.001), hip flexors (R: −0.397; p<0.001), knee extensors (R: −0.273; p=0.006), and knee flexors (R: −0.300; p=0.002) (Fig. 4A-D). For the upper limb, a significant negative correlation was observed for shoulder flexors (R: −0.405; p<0.001), shoulder abductors (R: −0.370; p<0.001), elbow flexors (R: - 0.337; p<0.001), and elbow extensors (R: −0.407; p<0.001) (Fig. 4E-H).

**Figure 4.**
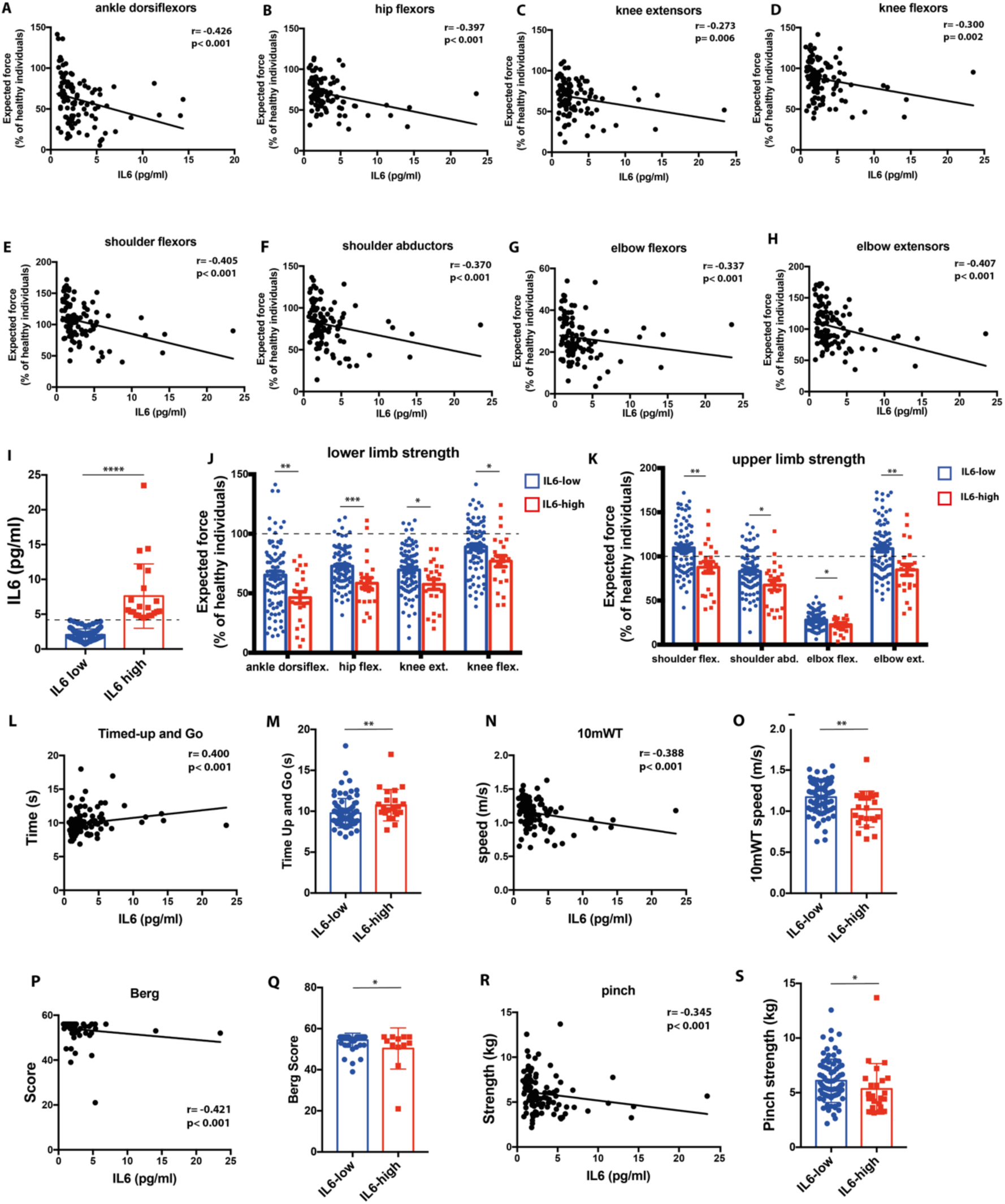
SASP expression is negatively correlated with muscle strength and functional outcomes in DM1. **A-D**) Correlation between serum IL-6 levels in DM1 patients and the expected strength (relative to normative values) of different muscle groups of the lower limb (ankle dorsiflexors, hip flexors, knee extensors, knee flexors) and **E-H**) upper limb (shoulder flexors, shoulder abductors, elbow flexors, elbow extensors). **I**) IL-6 expression levels in the blood serum of DM1 patients. Patients were classified in low- or high-expressing group based on the reference limit for healthy individuals (4.45 pg/ml, indicated by the dashed line). **J,K**) Expected muscle strength of DM1 patients with low or high levels of IL-6 for the lower limb (**J**) and upper limb (**K**) muscle groups. Dashed line indicates the normative data for isometric muscle strength in healthy individuals of the same age, sex, and weight. **L,N,P,R**) Correlation between serum IL-6 levels in DM1 patients and the results to different functional capacity tests: Timed-up and Go, 10-meter walk test (10mWT), Berg Balance scale, pinch test. **M,O,Q,S**) Functional capacity test results for DM1 patients with low or high levels of IL-6 levels. **I-K,M,O,Q,S**) Data are expressed as means +/-SEM. N=103 patients (80 patients IL-6 low, 23 patients IL-6 high). *p<0.05, **p<0.01, *** p<0.001.

Thereafter, we separated the group in patients expressing low or high levels of IL-6, based on the normative value for healthy individuals (upper 95th percentile reference limit of healthy individuals: 4.45 pg/ml)^32^. Twenty-three DM1 patients out of 103 (22.3%) had IL-6 levels over the reference limit of healthy individuals, suggesting that IL-6 is abnormally elevated in this population (Fig. 4I). The average IL-6 level was 3.7-fold higher (p<0.001) in the high IL-6 expressing group (7.61 pg/ml) compared to the low IL-6 expressing group (2.04 pg/ml). As expected, DM1 patients were weaker than normative values for healthy individuals for most muscle groups even if their IL-6 level was low (Fig. 4J,K). However, the DM1 patients that express higher levels of IL-6 are significantly weaker compared to those that had a normal SASP profile for all muscle groups tested: ankle dorsiflexors (−18%, p=0.006), hip flexors (−14%, p=0.001), knee extensors (−12%, p=0.015), knee flexors (−12%, p=0.021), shoulder flexors (−22%, p=0.006), shoulder abductors (−16%, p=0.016), elbow flexors (−5%, p=0.032), elbow extensors (−24%, p=0.003) (Figure 4J,K).

The functional capacity of DM1 participants was assessed using standardized physical tests validated in the DM1 population: the Timed-Up and Go (time to rise from a chair, walk 3 meters, walk back and sit down), 10-meter Walk-Test (10mWT; walking speed over a short distance), Berg balance scale (14-item test assessing static and dynamic balance), and pinch test (thumb to index pinch strength)^33, 34^. We observed a significant correlation between IL-6 levels and the scores at the Timed-Up and Go (R: 0.400; p<0.001), 10mWT (R: −0.388; p<0.001), Berg balance scale (R: −0.421; p<0.001), pinch test (R: −0.345; p<0.001) (Fig. 4L,N,P,R). There was a significant reduction in the group expressing higher levels of IL-6 compared to the one expressing lower levels of IL-6 in the performance at the different functional tests: Timed-up and Go (+1 s [+10%], p=0.005), 10 mWT (−0.15 m/s [-12%], p=0.001), Berg balance scale (−4.1 [-7%], p=0.025), and pinch test (−0.74 kg [-12%], p=0.032) (Fig. 4M,O,Q,S).

Further analysis using the stepwise regression model was used to verify if IL-6 is a candidate predictor variable explaining muscle strength and functional outcomes. Results show that IL-6, together with the clinical phenotype, age and/or sex of the patient, is a variable that explain the variance of the scores obtained by participants to the Timed-up and Go (standardized β: 0.18, p=0.048), 10mWT (standardized β: −0.2, p=0.003), Berg balance scale (standardized β: −0.3, p=0.01), and the predicted maximal muscle strength of hip flexors (standardized β: −0.27, p=0.02), knee extensors (standardized β: −0.20, p=0.034), knee flexors (standardized β: −0.19, p=0.034), ankle dorsiflexors (standardized β: −0.24, p=0.005), shoulder flexors (standardized β: −0.28, p=0.002), shoulder abductors (standardized β: −0.22, p=0.009), elbow flexors (standardized β: −0.18, p=0.034), and elbow extensors (standardized β: −0.206, p<0.001) (Suppl. table 4).

The same analyses were performed based on the serum level of TNFα. No correlation was observed between TNFα levels and the strength of ankle dorsiflexors, hip flexors, knee extensors, knee flexors, shoulder flexors, shoulder abductors, elbow flexors, elbow extensors, 10mWT, and pinch test (suppl. Fig. 7). Only the Berg balance test (R: 0.269; p=0.006) and the Timed-up and go (R: −0.305; p=0.009) showed a mild correlation with TNFα levels.

Altogether, our findings indicate that cellular senescence and SASP are hallmarks of DM1, which could explain the reduced myogenic capacity and the impaired muscle function. Therefore, we next explored the possibility of eliminating these senescent cells as a potential therapy to alleviate DM1 disease progression.

### Senolytics specifically eliminate senescent myoblasts in DM1 and reduce SASP expression

Considering that the pathophysiological mechanism behind cellular senescence varies depending on the inducers and cell types, we tested the impact of six different senolytics that act through different mechanisms to induce apoptosis of senescent cells: the FOXO4-DRI peptide (p53-FOXO4 interaction), Fisetin (PI3K/AKT pathway), Dasatinib + Quercetin (PI3K/AKT pathway), Navitoclax (BCL-2/BCL-X inhibitor), A1331852 and A1155463 (Selective BCL-X_L_ inhibitors)^24, 35–38^. Analysis of cell viability following the treatment of myoblasts with different concentrations of senolytics showed that FOXO4-DRI, Dasatinib + Quercetin, and fisetin were toxic for cells isolated from both healthy controls and DM1 patients (Suppl. Fig. 8). Navitoclax selectively eliminated cells in DM1 samples at a specific concentration (3 μm); however, A1155463 was the most effective to target cells specifically in the DM1 group and not affect the healthy cell viability (Fig. 5A). Moreover, A1155463 was effective at a lower concentration, in the nanomolar range. Therefore, this lead compound was chosen for the following experiments. To confirm that A1155463 induced apoptosis of senescent cells, an Annexin V/propidium iodide (PI) assay was performed on healthy and DM1 myoblasts. Treatment of myoblasts with A1155463 at a dose of 100 nM showed increased apoptosis only in the DM1 samples (Fig. 5B,C). SA-β -Gal staining on DM1 or controls myoblasts treated with vehicle or A1155463 showed a significant reduction in the number of SA-β -Gal+ senescent cells in the A1155463-treated DM1 group (Fig. 5D,E). Further analysis of senescence genes expression demonstrated that A1155463 treatment reduced *P16* expression in DM1 samples, but not in controls (Fig. 5F). The expression of SASP factors was analysed by multiplex Luminex assay in DM1 cells treated with A1155463 or vehicle. Results showed that senolytic treatment significantly reduced the expression of many SASP factors such as CSF3 (p=0.0005), CXCL1 (p=0.018), CXCL8 (p=0.002), CCL2 (p=0.041), MMP1 (p=0.009), and MMP3 (p=0.027) (Figure 5G). A trend toward reduction was also observed for IL-6 (p=0.078), CCL7 (p=0.054), MMP2 (p=0.06), and MMP12 (p=0.07). On the other hand, growth factors that stimulate cell proliferation such as EGF (p=0.0005) and FGF2 (p=0.026) were increased by A1155463 treatment (Suppl. Fig 9). Other anti-inflammatory cytokines and growth factors such as IL-4 (p=0.28), IL-13 (p=0.13), FLT-3L (p=0.25), and VEGF-A (p=0.38) were not significantly affected by the treatment (Suppl. Fig. 9).

**Figure 5:**
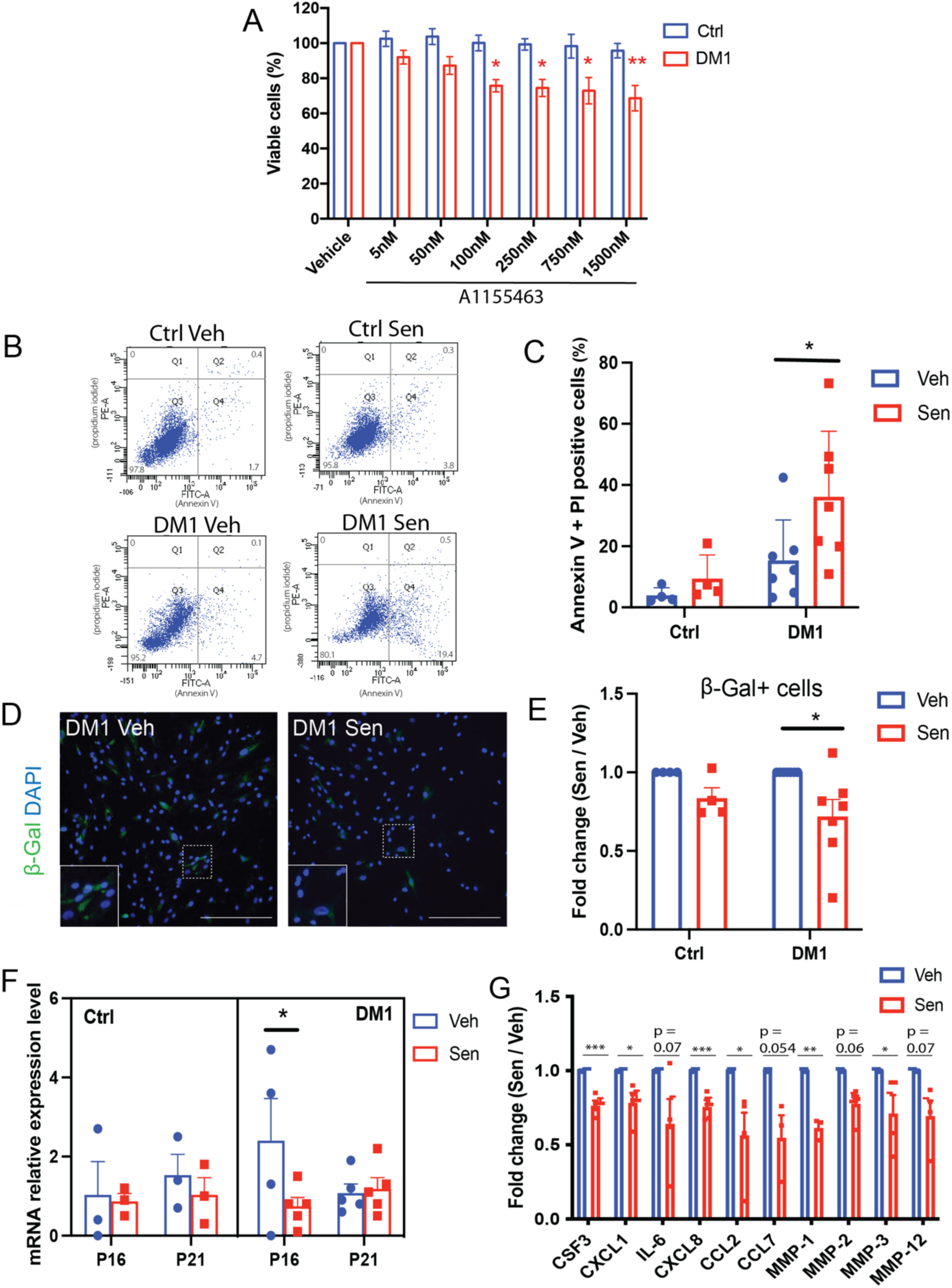
A1155463 shows senolytic activity on DM1 primary myoblasts culture. **A)** Cell viability assay of A1155463 at different concentrations using control (Ctrl) and DM1 patients’ myoblasts. *\*p= 0.038* (100 nM), *\*p=0.023* (250 nM), *\*p=0.011* (750 nM), *\*\*p=0.0016* (1500 nM). **B)** Representative FACS plot of Annexin-V (FITC) and propidium iodide (PE), and **C**) quantification of the percentage of positive apoptotic/necrotic cells after 48 h treatment of myoblasts (DM1 and Ctrl) with 100 nM of A1155463 or with vehicle; *\*p = 0.034.* **D)** Representative micrographs of DM1 myoblasts stained for the senescent marker SA-ß-Gal (green) after 72 h treatment with 100 nM of A1155463 or with vehicle. Scale bar: 300 μm. **E)** Quantification of the number of SA-ß-Gal+ cells expressed as fold change ratio of A1155463-treated (Sen) versus non-treated (Veh) cells (n = 4-7). *\*p = 0.015*. **F)** Quantitative real-time PCR for the senescence markers *P16* and *P21* on primary myoblasts after 72 h of treatment with vehicle or A1155463. *\*p = 0.042.* **G)** Multiplex luminex assay of SASP factors showing fold change ratio of A1155463-treated (Sen) versus non-treated (Veh) DM1 myoblasts (n = 3-4). *\*\*\*p= 0.0005* (CSF3), *\*p= 0.018* (CXCL1), *\*\*\*p= 0.002* (CXCL8), *\*p= 0.041* (CCL2), *\*\*p= 0.009* (MMP-1), *\*p=0.027* (MMP-3). Data are expressed as means +/-SEM.

### The eradication of senescent cells restores myogenesis in DM1

Next, we aimed to determine if the elimination of senescent cells, and the associated reduction in the expression of SASP factors, in A1155463-treated DM1 myoblasts restores the myogenic potential of the non-senescent cells. Myoblasts were treated with A1155463 or vehicle for 1 day and were allowed to recover in proliferating media for 3 days. Immunostaining for the proliferation marker KI67 revealed that A1155463-treatment increased the proportion of KI67+ cells in the DM1 samples (1.8-fold increase), but not in the controls (Fig. 6A-B). In another set of experiments, myoblasts were treated with A1155463 or vehicle for 1 day and incubated in low serum differentiating medium for 5 days. Immunofluorescence staining for the differentiation marker MYOG revealed a significant increase (2.2-fold) in the proportion of MYOG+ cells only in the DM1 samples, and not in the controls (Fig. 6C-D). This increase in MYOG expression in the DM1 samples was also confirmed by Western blot (Fig. 6E-F).

**Figure 6:**
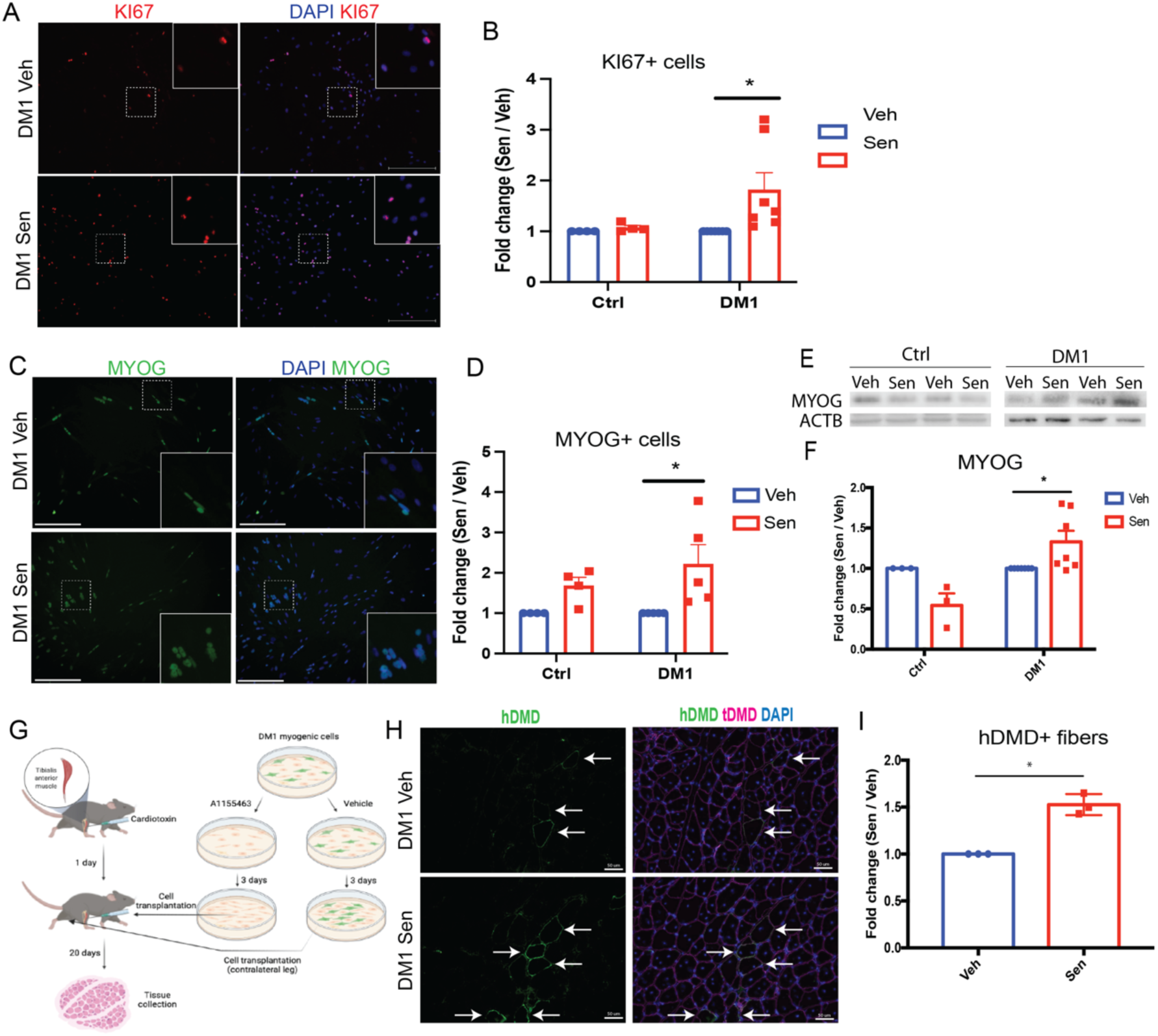
Clearance of senescent cells in DM1 improves myogenesis *in vitro* and *in vivo*. **A)** Representative micrographs of DM1 myoblasts immunolabeled with KI67 (proliferation marker, red) after 72 h of treatment with vehicle (Veh) or A1155463 (Sen). Scale bar: 300 μm. **B)** Quantification of the proportion of KI67+ cells expressed as fold change ratio of A1155463-treated (Sen) versus non-treated (Veh) cells (n = 4-7). *\*p = 0.017.* **C)** Representative micrographs of DM1 myoblasts differentiated for 5 days after treatment with vehicle (Veh) or A1155463 (Sen) and immunolabeled with MYOG (differentiation marker, green). Scale bar: 300 μm. **D)** Quantification of the number of MYOG+ cells expressed as fold change ratio of A1155463-treated (Sen) versus non-treated (Veh) cells (n = 4-5). *\*p = 0.0134.* **E)** Representative Western blot showing the expression of *MYOG* in control and DM1 myoblasts differentiated for 5 days after treatment with vehicle (Veh) or A1155463 (Sen). **F)** Quantification of MYOG expression by Western blot (relative to β-actin as loading control). Data are expressed as fold change ratio of A1155463-treated (Sen) versus non-treated (Veh) cells (n = 3-7). \**p= 0.039*. **G)** Schematic representation of DM1 myoblast transplantation experiment. **H)** Representative micrographs of muscle sections of TA muscles (21 days post-cardiotoxin injection) of NSG mice transplanted with DM1 myoblasts treated with vehicle (Veh, upper images) or A1155463 (Sen, bottom images). Sections were immunostained with human dystrophin (green), total dystrophin (red), and DAPI (nuclei, blue). White arrows indicate myofibers expressing human and total dystrophin. Scale bar: 50 μm. **I)** Quantification of the number of myofibers expressing human dystrophin expressed as fold change ratio of A1155463-treated (Sen) versus non-treated (Veh) cells (n = 3). *\*p = 0.014.* Data are expressed as means +/-SEM

To better address the *in vivo* myogenic potential of DM1 myoblasts treated with A1155163, we performed a cell transplantation assay. Three different DM1 myoblast lines were treated with A1155463 (100 nM for 24 h) or vehicle and cells were allowed to recover in proliferating media for 3 days. These cells were transplanted into the tibialis anterior muscle of NSG mice that were injured by cardiotoxin (CTX) injection 1 day before. Regenerated TA muscles were collected 21 days post-CTX injection (Fig. 6G). Cell engraftment was assessed by co-immunolabeling for the human dystrophin protein (hDMD) and total dystrophin protein (tDMD) (Fig. 6H). We observed that A1155463 enhanced DM1 cell engraftment by 50% when compared to vehicle (Fig. 6I). Taken together these results indicate that A1155463 treatment can significantly improve the myogenic potential of DM1 myoblasts *in vivo*.

## DISCUSSION

Our findings demonstrate that cellular senescence is a hallmark of myogenic cells in DM1. Using single cell transcriptomics and multiplex Luminex assay, we show that a subset of these cells displays a molecular signature characterized by high levels of expression of SASP factors. Our findings show that IL-6, a ubiquitous SASP factor, is associated with muscle weakness and functional limitations in DM1 patients. By screening for different senolytics, we identify a lead molecule, A1155463, that specifically eliminates senescent cells in DM1. The senolytic-induced reduction of SASP is associated with a restoration of the myogenic potential of DM1 myoblasts *in vitro* and *in vivo*. Altogether, these findings provide novel insights on the physiopathology of DM1 and identify a new therapeutic avenue for the treatment of this disease.

Signs of cellular senescence have been observed previously in cells isolated from skeletal muscles of DM1 patients; however, these studies used a mixed population of cells that were not purified by FACS and they did not validate if these senescence markers are co-expressed with myogenic cell markers^11–13^. Therefore, it was not possible to clearly delineate if myogenic cells, or if fibroblasts for instance, were becoming senescent. Particularly, fibroblasts or mesenchymal stromal cells are a predominant senescent cell type in various pathogenic conditions^39^. Here, we use FACS purified myogenic cells and we have confirmed that senescence markers (e.g., P16) are expressed in DM1 purified myoblasts. Moreover, co-staining for P16 positive cells and PAX7 on skeletal muscle biopsies revealed the presence of senescent muscle stem cells *in situ*, which has never been described before. These findings indicate that cellular senescence in DM1 is not only an *in vitro* artefact caused by replicative stress, but a relevant pathophysiological process of the disease. Furthermore, our experiments were performed on samples from individuals with the infantile, juvenile or adult form of the disease, while previous studies focused exclusively on the more severe congenital form, suggesting that cellular senescence is a hallmark of the disease no matter the clinical form^12, 40^.

Our findings show that the accumulation of RNA foci in the nuclei of DM myoblasts is associated with cellular senescence, suggesting that RNA-meditated toxicity is the cause of senescence in this disease. Toxic RNA accumulation caused by the CTG repeats in the *DMPK* gene is known to induce reactive oxygen species (ROS) production and oxidative damage in DM1^14, 41–44^. Chronic ROS production triggers a DNA damage response leading to cell cycle arrest and cellular senescence, which in turn secretes further ROS and SASP leading to a feed-forward cycle^14, 45, 46^. Using C2C12 cells it was shown that the insertion of CUG repeats increases ROS production, which is further exacerbated by H_2_O_2_ exposure, suggesting that DM1 cells produce more ROS and are more sensitive to oxidative stress^14^. Analysis of the GO enrichment data of our single cell transcriptomic dataset show that the most dysregulated genes in DM1 patients’ cells are involved in oxidative phosphorylation. Moreover, pathway analysis also shows an enrichment for genes related to ROS and reactive nitrogen species (RNS) production. Analyses from other datasets also show an increase in ROS producing genes (e.g., *NOX4*) and a decrease in antioxidant genes (e.g. *SOD3*)^14, 47^. Notably, the expression of antioxidant genes such as *SOD1*, *SOD2*, and *GPX1* have been positively associated with muscle strength in another DM1 transcriptomic dataset, suggesting that antioxidant levels could help protect against senescence-associated muscle wasting^48^. Overall, our findings and others suggest that senescence in DM1 myoblasts is induced by toxic RNA-mediated oxidative stress.

Other transcriptomics datasets have been generated on DM1 tissues and cells (reviewed in ^49^). A study on DM1 myogenic cells showed that the transcriptional changes are more important in DM1 myoblasts than myotubes^47^. GO term analysis showed that the biological processes that are mostly enriched in DM1 myoblasts are signalling pathways, ECM components, and cytoskeletal organisation^47^. Another RNAseq dataset performed on human embryonic stem cells (DM1 and controls) differentiated into myogenic cells showed that inflammatory pathways such as IL6-JAK-STAT3 signalling are overexpressed in DM1 cells, which is coherent with the inflammatory signature that we observed^50^. Similarly, RNAseq analysis performed on isogenic myoblasts containing or not a CTG_2600_ expansion showed an enrichment in biological processes such as cellular response to IFN*γ*^51^. Using a model of C2C12 cells containing expanded CUG repeats, it was also shown that toxic RNA expression induces a senescence-like signature characterized by enrichment in genes related to ECM organisation, response to oxygen levels, apoptosis, and regulation of cell growth^14^. Transcriptomic analysis on skeletal muscle samples from individuals with the congenital form of DM1 showed a signature very similar to our scRNAseq dataset characterized by downregulation in myogenesis genes and upregulation in inflammatory genes, such as the IL6/STAT3 target genes *SAA1* and *SAA2*^40^. Moreover, supporting our results, RNAseq on muscle samples from DM1 patients and controls showed that genes related to SASP factors (e.g. *CXCL14, CXCL16*, and *MMP2*) or cell cycle inhibitors (e.g. *CDKN1A*, *TP63*)^52^, are negatively correlated with muscle strength in DM1^48^. RNAseq performed in other tissues such as the frontal cortex showed an enrichment in inflammatory genes compared to healthy individuals, suggesting that the pro-inflammatory signature is a common feature in different DM1 tissues^53^. These findings bring important insights on the molecular mechanisms in DM1, however, bulk RNAseq analysis limits the interpretations regarding the subpopulation of cells affected. Particularly, it is not possible with these datasets to determine if the inflammatory signature is generalized or circumscribed to a specific subset of cells, which is especially important in the context of cellular senescence.

To assess the specific molecular signature of myogenic cells in DM1, we used an unbiased approach through scRNAseq. This is the first single cell transcriptomics dataset generated in DM1^49^. This dataset allowed to identify specific cell clusters expressing different levels of senescence markers leading to a better comprehension of the different stages of senescence in myoblasts of patients with DM1. Particularly, we identified a specific cluster of DM1 myoblasts expressing high levels of SASP factors, while another cluster expressed higher levels of *SAA1* and *SAA2*, two molecules that are triggered by SASP factors and can further stimulate SASP expression by a positive feedback loop^27, 54^. These findings suggest that the expression of SASP by senescent cells have paracrine effects on neighbouring cells and promote their senescence-like phenotype. Moreover, this dataset identified that, while a subset of cells shows a fully senescent or early senescent profile, there is a significant proportion of cells that had a similar molecular profile as the one found in our control samples from healthy individuals. These results suggest that elimination of the senescent myoblasts could restore the myogenic activity of non-senescent DM1 myoblasts. Several studies used this approach to alleviate chronic and degenerative diseases in which senescence contributes to the progression of the disease (reviewed in ^55, 56^). By testing different senolytic drugs, we observed that DM1 senescent myoblasts were particularly responsive to inhibitors targeting the BCL family members. Navitoclax (ABT-263; BCL2/BCL-XL inhibitor) was effective at targeting senescent myoblasts in DM1, similar to what was shown in an aging model^38^; however, the highest efficiency was achieved with the BCL-XL inhibitor A1155463. Notably, our SCENIC analysis showed an enrichment in the activity of the transcription factors of the NF-kB family, which are known to bind to promoter region of Bcl-xl gene to stimulate its expression^57^. Other widely used senolytic such as FOXO4-DRI, fisetin, or dasatinib + quercetin that affect other pathways were ineffective to specifically target senescent DM1 myoblasts^24, 36^, which indicate that the senescent molecular signature expressed in DM1 myoblasts is distinct to this cell type/inducer and that specific molecular targeting is necessary to eliminate these cells.

Our findings demonstrate that IL-6, a main component of the SASP cocktail, is correlated with muscle weakness and functional capacity limitations in DM1. These results are consistent with previous observations showing that members of the IL-6 signalling pathway are correlated with signs of histological degenerative changes in the congenital form of the disease^40^. On the other hand, our findings show that another pro-inflammatory cytokine, TNFα, was not correlated with muscle weakness. These results are consistent with our scRNAseq data showing an increase in the expression of *IL6* but not *TNF* in samples from DM1 patients. These results suggest that there is a specific SASP signature associated with muscle weakness in DM1, and not only a generalized pro-inflammatory profile. Similarly, another study screened for 20 pro-inflammatory cytokines in the serum of patients affected by facioscapulohumeral muscular dystrophy and identified IL-6 as the only cytokine correlating with muscle weakness and disease severity^58^. These findings suggest that IL-6 is a promising biomarker to determine the severity of muscle dysfunction in DM1^59^.

Our results also suggest that targeting SASP cytokines such as IL-6 is a valid therapeutic approach to mitigate disease progression. Our findings showed that specific elimination of senescent DM1 myoblasts leads to a reduction in the expression of SASP molecules, which restores the myogenic function of non-senescent DM1 myoblasts through an improvement of their cellular proliferation and differentiation capacity. These results are consistent with a previous study showing that the secretion of SASP by senescent cells block myoblast differentiation *in vitro*^60^. The improvement in myogenesis was not observed in cells from healthy individuals, which confirms that the senolytic drug targets a specific pathophysiological mechanism in DM1.

Overall, this study describes a well-defined senescent molecular signature in DM1 myogenic cells, which brings novel insights on the pathophysiological mechanisms of the disease. Particularly, it identifies the production of SASP as a major characteristic of DM1 myoblasts, which is associated with muscle weakness in DM1. We demonstrated that specific drugs can be used to target these defective cells and restore myogenesis. These findings open a new therapeutic avenue that could help to mitigate the impact of DM1, a disease for which no treatment currently exists to slow symptoms progression^61^.

## METHODS

### Participants’ recruitment

Patients were recruited at the Saguenay Neuromuscular Clinic of Jonquière (Québec, Canada) and healthy individuals were recruited among the friends/family members or team members. Inclusion criteria for DM1 participants were to be aged over 18-year-old and have a genetically confirmed diagnosis of DM1. One hundred and three DM1 participants were recruited between 2002-2004 and completed muscle and functional capacity assessments and blood sample collection. Ten DM1 patients and seven healthy individuals were recruited in 2019 and undergone a muscle biopsy only. This study was approved by the Ethics review board of the Centre de santé et de services sociaux de Chicoutimi, Canada. Written informed consent was obtained from all participants.

### Muscle strength, functional capacity assessments, and blood sample collection

A blood sample was taken from each participant and the level of IL-6 was measured at the *Centre de recherche sur les maladies lipidiques* of the *Centre hospitalier de l’Université Laval* (Québec, Canada). DM1 patients were classified in a low or high IL-6 expressing group based on the cut-off value representing the upper 95th percentile value for healthy individuals (4.45 pg/ml)^32^. The maximum isometric muscle strength of the ankle dorsiflexors, hip flexors, knee flexors, knee extensors, shoulder flexors, shoulder abductors, elbow flexors and elbow extensors was assessed using quantitative muscle testing according to a standardized protocol^62^. Muscle strength was expressed as the expected muscle strength of healthy individuals based on a French isometric strength normative database using predictive equations taking into consideration age, sex, and weight of the patients^31^. Patients were subjected to a battery of validated functional tests^63, 64^. Timed-up and Go: time to stand up, walk three meters, turn and walk back to sit down on the chair; 10mWT: comfortable walking speed (m/s) along a 10-meter distance; Berg balance scale: 14 items measuring static and dynamic balance (maximum score of 56 indicating no deficit in balance); Pinch strength: measured using a B&L pinch gauge (kg). All tests were conducted under the supervision of a trained physiotherapist

### Muscle biopsy

A biopsy of the vastus lateralis at the mid-thigh level was taken using the modified Bergstrom needle technique with suction^65^. A ∼200 mg sample of muscle was obtained, rinsed in sterile phosphate buffered saline solution (PBS) and portioned out for the various following analyses. The first half of the sample (approximately 100 mg) was kept in sterile SK-MAX media (Wisent Bio) on ice for muscle stem cell isolation. The second half of the muscle sample was separated in two parts that were flash frozen for qPCR analysis (50 mg) or embedded in OCT tissue freezing medium and frozen in 2-methylbutaned cooled in liquid nitrogen (50 mg). The samples were stored at −80°C.

### Myogenic cell lines isolation, purification, and culture

Muscle biopsies from controls and patients were minced, digested in collagenase, and plated in a culture dish^66^. Myoblasts were grown in Sk-MAX media complemented with Sk-MAX supplement and 20% fetal bovine serum (FBS). Myoblasts were expanded for a few days until there were enough cells for purification by FACS. Myoblasts were stained with 7-AAD (cell viability) and with the positive selection marker anti-CD56 AF647-conjugated (clone R19-760, BD biosciences). Of note, preliminary experiments showed that virtually all CD56+ cells were also positive for another myogenic marker CD82 (data not shown) ^67^, and therefore only CD56 was used for the follow-up experiments. Myoblasts differentiation was induced by exposing cells to 2% horse serum (HS) in DMEM media. Control and DM1 myoblasts were used for experiments at the same low passage (maximum P6).

### Single-cell RNA-sequencing experiments

Single cell suspensions of myogenic cells were resuspended in PBS containing 0.04% ultra-pure BSA. scRNAseq libraries were prepared using the Chromium Single Cell 30 Reagent Kits v3.1 (10x Genomics; Pleasanton, CA, USA) according to the manufacturer’s instructions^68^. Generated libraries were sequenced on an Illumina HiSeq4000.

Raw sequencing data were preprocessed using Cell Ranger 6.1.2^69^ to perform the alignment (human genome GRCh38-2020-A), filtering, barcode counting, and UMI counting at a single cell resolution, generating a control, patient, and a merged digital gene expression (DGE) matrix containing 1,902, 2,029 and 3,931 cells respectively. Further downstream analyses were conducted using R package Seurat v4.0 (spatial reconstruction of single-cell gene expression data^70^) and singleCellTK v2.2.0 (Single cell Toolkit).

Quality control: Due to deep sequencing of the libraries, for each DGE matrix, cells expressing fewer than 1,000 genes and more than 10% of mitochondrial genes were filtered out, leaving a total of 3,381 cells in the merged matrix (1,639 cells from controls and 1,742 cells from patients). For all cells, genes were excluded if they were expressed in less than 3 cells. We normalized the DGE matrices by the total number of UMIs in each cell and scaled by 10^6^ to yield counts per million (CPM). We natural log transformed these matrices after the addition of a pseudocount of 1 to avoid undefined values.

The 2,000 most variable genes were selected for each DGE matrix, based on their relative dispersion (variance/mean) regarding to the expected value across genes with similar average expression^71^. Next, we performed a linear transformation (‘scaling’) to each matrix to minimize variations due to technical noise or possible batch effects.

We used DoubletFinder^72^ to identify putative doublets in each DGE matrix relying solely on gene expression data. DoubletFinder requires three input parameters for the doublet classification: the number of expected real doublets (nExp), the number of artificial doublets (pN) and the neighborhood size (pK). We calculated the nExp by multiplying the number of cells by the expected doublet ratio (1.6%) based on 10x genomics Chromium user guide. To identify pN and pK values, we performed the parameter optimization of DoubletFinder resulting in a pN of 0.30 for all matrices and a pK of 0.24, 0.01 and 0.005 for the control, patient, and merged matrices respectively. After the removal of putative doublets, we remain with 1,613 cells in the control, 1,714 cells in the patient, and 3,327 cells in the merged matrix for downstream analyses.

We estimated cell cycle activity by scoring the expression of a set of S phase– associated and G2/M phase–associated genes, as implemented in Seurat^70^.

To enable unsupervised clustering, we performed a linear dimensionality reduction with principal component analysis (PCA) on each DGE matrix using the highly variable genes previously identified. Once embedded in this PCA space, we applied a K-nearest neighbor (KNN) graph-based clustering approach and refine the edge weights between any two cells based on the shared overlap in their local neighborhoods, identifying the k = 10 nearest neighbors for each cell. To cluster the cells, we apply a Louvain algorithm^73^ to iteratively group cells together, with the goal of optimizing the standard modularity function. A resolution of 0.3 was used in the patient matrix to identify more defined cell populations. The t-Distributed Stochastic Neighbor Embedding (tSNE)^74^ and the Uniform Manifold Approximation and Projection (UMAP) were used to visualize the clustering results; however, UMAP was chosen because it provides a better separation of the groups.

We computed differentially expressed genes (DEGs) between controls and patients using the Wilcoxon rank-sum test with the Benjamini-Hochberg procedure for FDR control^75^, identifying positive and negative markers for each group. DEGs with |logFC| > 1.0 and adjusted p-values < 10e-6 were considered. In addition, we also computed the proportion and the probability distribution of cells expressing each gene in each group. To identify DEGs among patient clusters and patient cells after the senescence annotation (method described below), we used the model-based analysis of single-cell transcriptomics method (MAST)^76^. DEGs with |logFC| >0.25, FDR <0.05, percentage cluster expression > 0.5, percentage expression in the control group < 0.4, and adjusted p-values <0.01 were considered. For the senescent and non-senescent populations, DEGs with an adjusted p-values <0.01 and logFC > 1.5 in one condition and logFC <1.5 in the other condition were plotted regardless the expression of the control.

Gene Ontology terms were analyzed for DEGs identified in each cell type using the online Metascape tool (https://metascape.org)^77^ with default parameters: Min Overlap = 3, p-value Cut-off = 0.05, and Min Enrichment = 1.5. The top 20 terms ranked by p-value were displayed in a bar diagram. Pathway enrichment analysis was performed using R package ReactomePA v1.36.0 and displayed in a bubble diagram^78^.

To perform the annotation of senescent and non-senescent cells in patient cells, we used the R package GSVA v1.40.1 ^79^ to compare the average expression of the DEGs of each patient cluster with reference gene sets related to cellular senescence and secretory processes^28, 80–82^. Heatmap and density plots are displayed. Clusters with GSVA scores > 0 in all gene sets were annotated as fully senescent cells, clusters with GSVA scores < 0 in all gene sets were annotated as non-senescent cells, and clusters with positive and negative GRVA values were annotated as early senescent cells.

### Senescence associated beta-galactosidase staining

CellEvent, senescence green detection kit (ThermoFisher, C10850) was used to label senescent cells according to the manufacturer’s specifications. The fluorescent probe included in the assay is a substrate for beta-galactosidase and emits a 488 AlexaFluor fluorogenic signal in cells containing the beta-galactosidase active enzyme.

### Fluorescence in situ hybridation (FISH)

Following the SA-β-gal staining protocol, cells were fixed with 2% PFA solution for 10 min at room temperature, washed, and dehydrated in prechilled 70% ethanol for 3 h at 4°C. Thereafter, cells were washed in PBS RNAse free for 10 min, incubated in 40% formamide (Millipore, Ca) in 2× saline-sodium citrate (SCC) buffer (300 mM sodium chloride, 30 mM sodium citrate, pH 7.0) for 10 min at room temperature, and blocked in hybridization buffer (40% formamide, 2× SCC buffer, 200μg/mL bovine serum albumin, 100 mg/mL dextran sulfate, 2 mM vanadyl sulfate, 1 mg/mL yeast tRNA) for 15 min at 37°C^83^. The Cy5-labeled (CAG)10 DNA probe was denatured for 10 min at 100°C, chilled on ice for 10 min, then added to prechilled hybridization buffer for a final concentration of 150 ng/μL probe. Cells were incubated with the probe for 2 h at 37°C. As a negative control, RNase solution were incubated for 10 min at RT before the hybridization of the probe. Cells were washed, counterstained with DAPI, and images were acquired on SP8 confocal microscope (Leica).

### RT-qPCR

Myogenic cells total RNA was extracted with Qiazol reagent according to the manufacturer’s specifications. Total RNA was quantified with a Thermo Scientific™ NanoDrop™ 8000 Spectrophotometer. Reverse transcription was performed on 1 μg of total RNA using the RT mastermix (QuantiTect Reverse Transcription Kit, 205313) to obtain cDNA. qPCR was performed with a set of primers designed on Primer-BLAST (NCBI) and validated for their specificity, efficiency and annealing temperature^84^. Gene amplification was performed with the BlasTaq™ 2X qPCR MasterMix (Abm, G892) on a Roche LightCycler® 480 Instrument II. Data were analysed with the LightCycler® 480 and were normalised relative to RPLPO expression^85^. The primers used are shown in Supplemental Table 5.

### Immunofluorescence

Muscle biopsies were cut at 10 μm thick in using a NX50 Cryostar (Thermo). Slices were put on positively charged Superfrost slides. Immunostaining was performed on skeletal muscle sections or on primary myoblast cultured on plastic dishes. Samples were fixed with 2% PFA for 5 min (sections) or 10 min (cells). Sections were incubated with a blocking solution containing 5% of donkey serum and 2% of bovine serum albumin (BSA) in PBS for 60 minutes at room temperature. Sections were incubated overnight at 4°C with the following primary antibodies diluted in blocking solution: PAX7 (clone PAX7, 1:20; Developmental Studies Hybridoma Bank, DSHB), P16 (ab54210, 1:50; Abcam), P21 (clone F5, sc-6246, 1:50; Santa Cruz), KI67 (14-5698-82, 1:500; Invitrogen), MYOG (ab124800, 1:200; Abcam), hDMD (MANDYS106(2c6), 1:2; Developmental Studies Hybridoma Bank, DSHB), total DMD (ab15277, 1:200; Abcam). Secondary antibodies were also diluted in blocking buffer (1:1,000) and incubated for 1 h at room temperature. Secondary antibodies (Invitrogen) used were donkey anti-mouse 488 (A21202) or donkey anti-mouse 594 (A21203), donkey anti-rabbit 488 (A21206) or donkey anti-rabbit 647 (A32795). Slides were mounted using PermaFluor mounting medium (Fisher Scientific). Immunofluorescence pictures of samples were taken with EVOS M5000 (Thermo Fisher Scientific) or Leica TCS SP8 DLS confocal microscope. Pictures were analyzed using ImageJ software (version 1.53, National Institutes of Health, USA).

### Viability assay

Cell viability was determined by Cell-titer 96 aqueous one solution cell proliferation assay kit (Promega, G3580) according to the manufacturer’s specifications. Briefly, aliquots of 5 × 10^3^ cells/well were cultured in 96-well plates. After 24 hours in culture, cells were treated with vehicle (DMSO) or the following senolytics: FOXO4-DRI (NovoPro), Fisetin (Selleck chemicals), Dasatinib + Quercetin (Sigma), Navitoclax (ABT-263, Abcam), A1331852 (Selleck Chemicals), or A1155463 (Selleck Chemicals), at different concentrations for 48 h. Then, 40 μl of Cell-titer 96 aqueous one solution was added to each well and incubated for an additional 3 h. The absorbance at 490 nm was recorded with a 96-well plate reader (BMG Labtech, CLARIOstar).

### Apoptosis assay

Apoptotic cells were identified using a detection kit (Biolegend, FITC Annexin V Apoptosis Detection Kit with PI, 640914) according to manufacturer’s specifications. After trypsinization, cells were washed twice with Cell Staining Buffer and stained with FITC Annexin V (1:30) and PI (1:10) for 15 minutes at room temperature. Then, 400 μl of Annexin V binding buffer was added, and the samples were analyzed by LSR Fortessa cytometer (BD Biosciences).

### SASP markers Luminex assay

A multiplex Luminex assay (Eve Technologies, Calgary, Alberta) was used to measure SASP markers of myogenic cells supernatant media. Two different discovery assays were chosen to assess SASP markers: Human Cytokine/Chemokine 48-Plex Discovery Assay® Array (HD48) and Human MMP and TIMP Discovery Assay® Array. Cells were treated with senolytic or vehicle and incubated for 36 hours. Cell media was changed, and cells were incubated for another 36 hours before supernatant collection. Supernatants were stored at −80 °C until assayed.

### Western blot

Cells were washed with sterile PBS and lysed with ice-cold RIPA buffer containing 1% of protease inhibitors and centrifuged at 10,000 g for 10 min. The supernatant was retained, aliquoted, and the protein content was quantified using the BCA Assay Kit (Thermo scientific, Mississauga, Ontario, Canada). A volume corresponding to 40 μg of protein was diluted with a sample buffer (125 mM Tris buffer (pH 6.8), 4% SDS, 20% glycerol, 0.05% bromophenol blue, and 200mM dithiothreitol), heated at 100°C for 5 min and electroseparated on 15% sodium dodecyl sulfate-polyacrylamide gel. Proteins were transferred to polyvinylidene difluoride membranes, which were blocked with 5% non-fat milk or 5% BSA for 90 min at room temperature. Membranes were immunoblotted overnight at 4°C with anti-myogenin (ab124800, 1:500; Abcam) and anti-beta-actin (4967S, 1:5,000; Cell Signaling) as primary antibodies. After washing, membranes were incubated with goat anti-rabbit (H+L) HRP-conjugated secondary antibodies (1:5,000; Abcam, ab6721) for 1 h at room temperature. Bands were revealed with ECL-plus Western blotting reagent (PerkinElmer Life and Analytical Sciences, USA), visualized with the BioRad ChemiDoc Imaging System, and quantified using ImageJ (National Institutes of Health, Maryland, USA). Densities were normalized to the loading control.

### Cell transplantation assay

NOD/SCID/IL2Rγ null (NSG) mice were housed in the animal care facility at the CHU Sainte-Justine Research Center under pathogen-free conditions in sterile ventilated racks. All *in vivo* manipulations were previously approved by the institutional committee for good laboratory practices for animal research (protocol #2020-2357). To induce muscle regeneration and enhance cellular engraftment, a cardiotoxin injection (Latoxan, 50 μl of 10 μM solution in saline) was performed intramuscularly (i.m.) through the skin in the right and left tibialis anterior (TA) muscles under general anesthesia. After 24 hours, 2 × 10^5^ DM1 myoblasts (resuspended in 10 μl of PBS) that were previously treated with A1155463 or vehicle for 72 hours were transplanted into the right or left injured TA muscle, respectively. Three different cell lines were used, and each cell line was transplanted into 3 different mice. Mice were allowed to recover for 20 days after which they were euthanized by CO_2_ inhalation (under anesthesia) followed by cervical dislocation and their TA muscles were collected. Tissues were embedded in OCT tissue freezing medium and frozen in 2-methylbutaned cooled in liquid nitrogen and stored at −80°C until sectioning.

### Statistical analysis

Sample size determination was based on the expected effect size and variability that was previously published for similar readouts using DM1 patient myogenic cultures^5,12^. No data were excluded. All experiments were repeated independently at least twice in the laboratory with similar results. For each set of experiments, the proportion of control and DM1 patients were matched as best as possible for age and sex. For data collection and analysis, the experimenter was blinded to the identity of the sample. For *in vitro* experiments, data were analyzed with the MIXED procedure of the Statistical Analysis System (SAS Institute, version 9.2, Cary, North Carolina, USA). Normality was verified for all data according to the Shapiro–Wilk test. Treatment effects were determined with two-tailed Student’s t-tests, One-way or Two-way analysis of variance (ANOVA) uncorrected Fisher’s Least Significant Difference (LSD) test. Results are reported as mean ± standard error of the mean (SEM). For i*n vivo* data analyses, associations between IL-6 level and muscle strength and functional capacity of patients were assessed using Spearman ρ correlation coefficient. Comparisons of muscle strength and functional capacity between patients based on IL-6 normative value for healthy individuals (below or over the threshold) were performed using Mann-Whitney U Test. Stepwise multiple regression analyses were used to determine which variables among age, sex, phenotype and IL-6 level (low or high) best explain each variable of muscle strength and functional capacity. For all tests, the significance value was set at *p*-value <.05. Data were analyzed using IBM SPSS Statistics for Windows, Version 24.0 (Armonk, NY: IBM Corp).

## ACKNOWLEDGEMENTS

We thank M. M. Hicham Affia, Ms. Veronique Lisi, and Ms. Lydia Tellier for their technical assistance. T.C.C. was supported by a fellowship from the Myotonic Dystrophy Foundation. G.D.B. was supported by CHU Sainte-Justine, University of Montreal, and RQR (réseau québécois en reproduction) awards. Z.O. was supported by an award from the MITACS fellowship. I.M. was supported by CERMO-FC (Center of Excellence in Research on Orphan Diseases – Fondation Courtois) and TheCell awards. A.D. was supported by an FQRNT (Fonds Québécois de la recherche - nature et les technologies) doctoral award. M.P.R. was supported by an FRQS (Fonds Québécois de la recherche – Santé) doctoral award. G.A. holds the Banque Nationale Research Exellence Chair in Cardiovascular Genetics. E.D. was supported by FRQS Junior-1 award and N.A.D. and S.M. were supported by FRQS Junior-2 award. This study was supported by grants from Canadian Institutes of Health Research (CIHR) (grant no. MOP-49556, JNM-108412), the Quebec Cell, Tissue and Gene Therapy Network –ThéCell (a thematic network supported by the FRQS) and Muscular Dystrophy Canada.

## AUTHOR CONTRIBUTIONS

N.A.D. and E.D. conceived and managed the project. T.C.C, G.D.B, Z.O., I.M., M.P.R, A.D., O.P., D.M., I.C., B.B., S.L., J.M., C.G. designed and performed experiments, and collected data. J.M., C.G., G.A., C.B., S.M, T.C.C, L.F., G.D.B, I.C., E.D. and N.A.D. analyzed and interpreted the data. T.C.C, G.D.B, E.D., and N.A.D wrote the manuscript. All authors carefully reviewed and provided critical insights to the manuscript.

