## supplemental material for "Clearance of defective muscle stem cells by senolytics reduces the expression of senescence-associated secretory phenotype and restores myogenesis in myotonic dystrophy type 1"

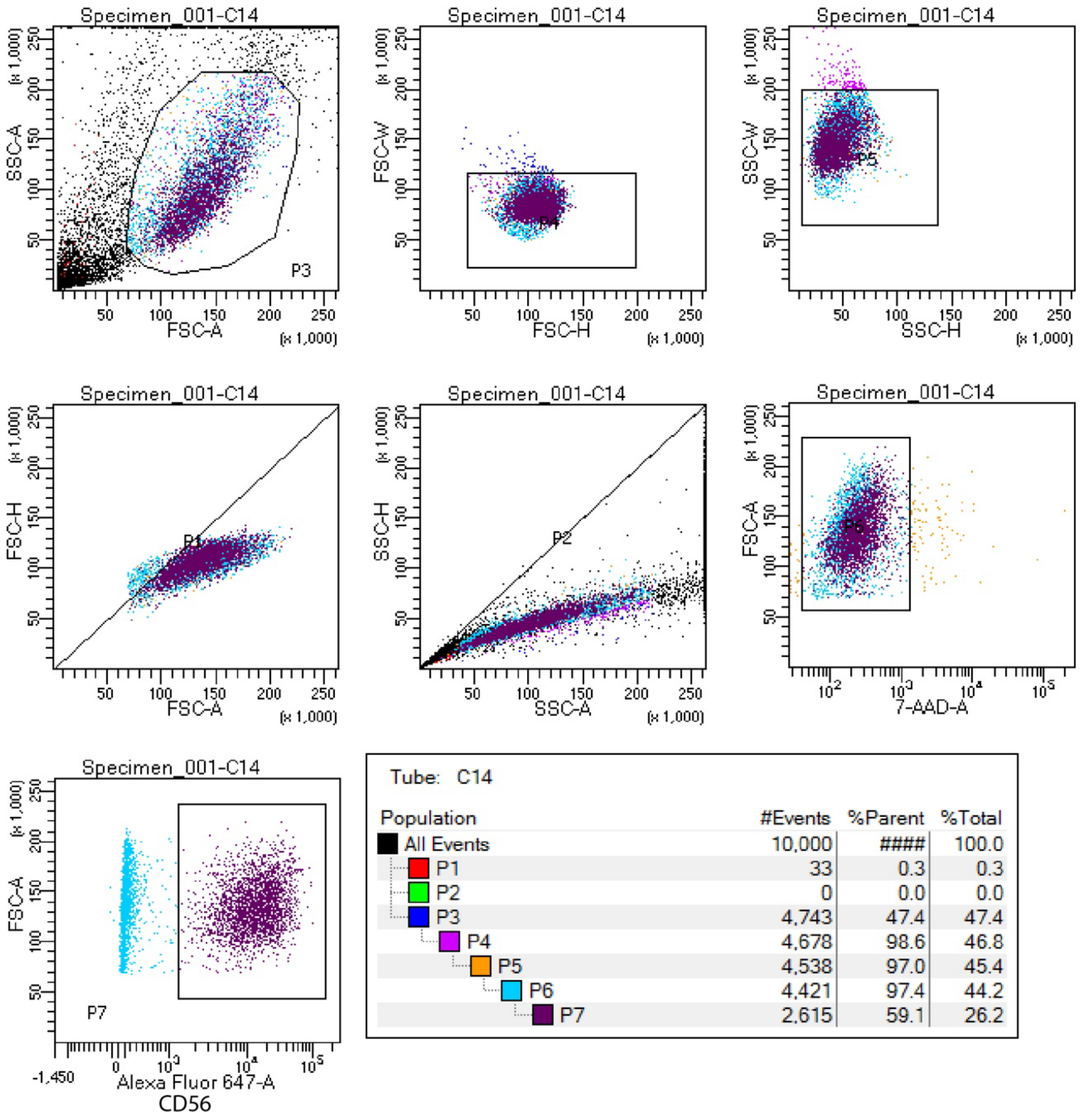


**Supplemental Figure 1. FACS gating strategy**. Representative FACS plot showing the gating strategy to purify human myogenic cells using forward scatter (FSC), side scatter (SSC), 7AAD (cell viability), and CD56 AF647-conjugated antibody (human myogenic cell marker).

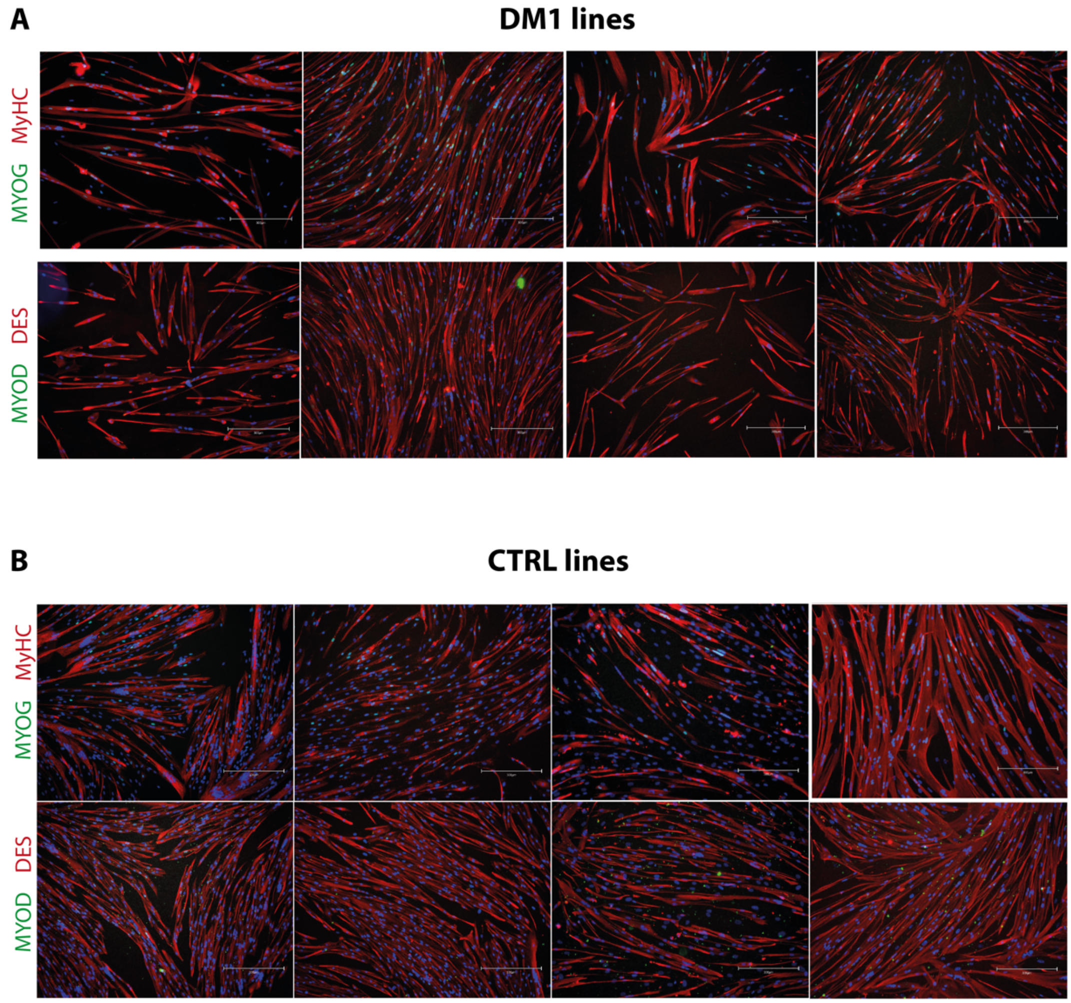


**Supplemental Figure 2. Myogenic cell purity.** Representative micrographs of immunofluorescent labelling of the myogenic markers Myogenin (Myog, green) and Myosin heavy chain (MyHC, red), or desmin (DES, red) and MYOD (green) on **A**) DM1 myoblast and **B**) healthy control cell lines.

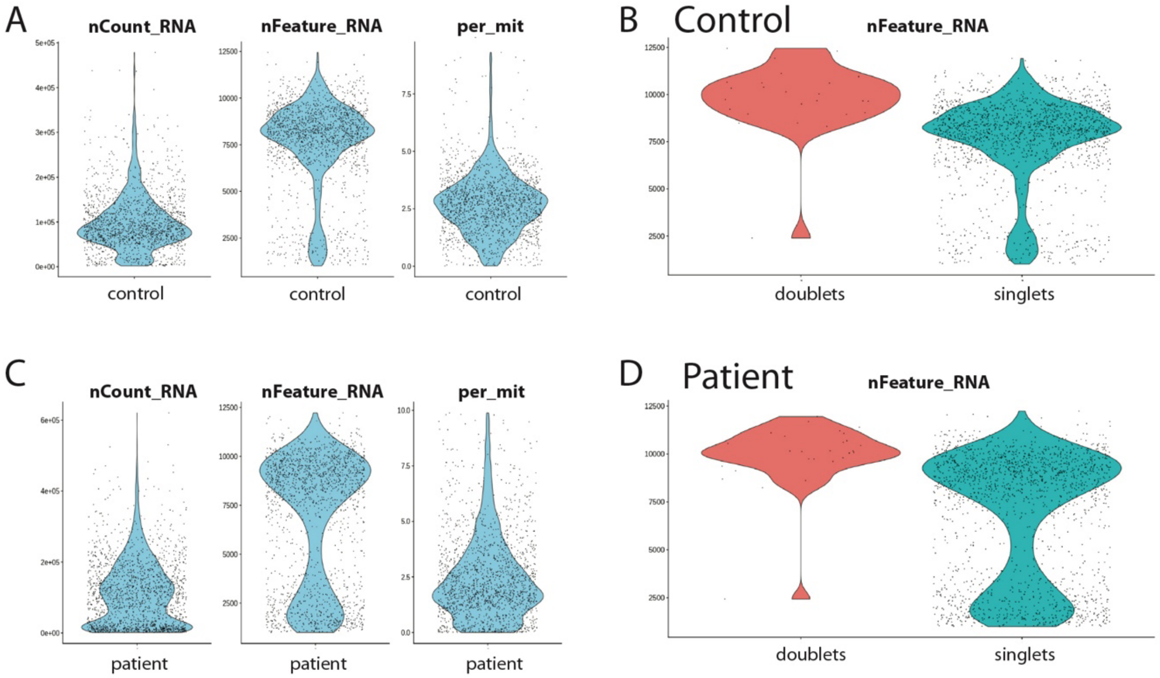


**Supplemental Figure 3. scRNAseq quality control. A,C)** Violin plots showing quality control features (nCount RNA, nFeature RNA, percent mito) and **B,D)** identification of singlets vs doublets for healthy controls (**A,B**) and DM1 myoblasts (**C,D**) scRNAseq analysis.

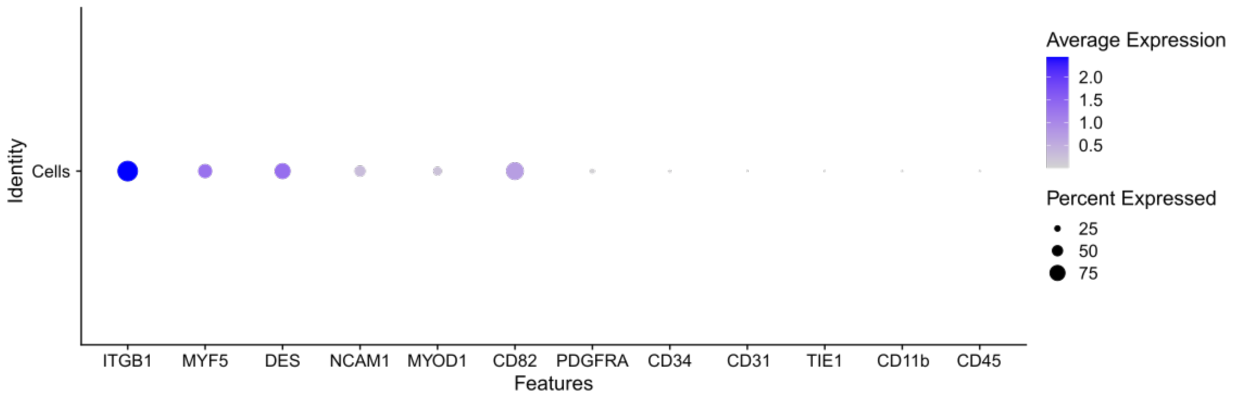


**Supplemental Figure 4. Cell type markers in scRNAseq analysis.** Dot plot showing the percentage of cells (size of the dots) and the expression level (color of the dots) of myogenic genes (*ITGB1, MYF5, DES, NCAM1, MYOD1, CD82*) and non-myogenic genes: fibroadipogenic progenitors (*PDGFRA, CD34*), endothelial cells (*CD31, TIE1*), and myeloid cells (*CD11B, CD45*) from the single-cell RNAseq dataset of DM1 patients and healthy controls.

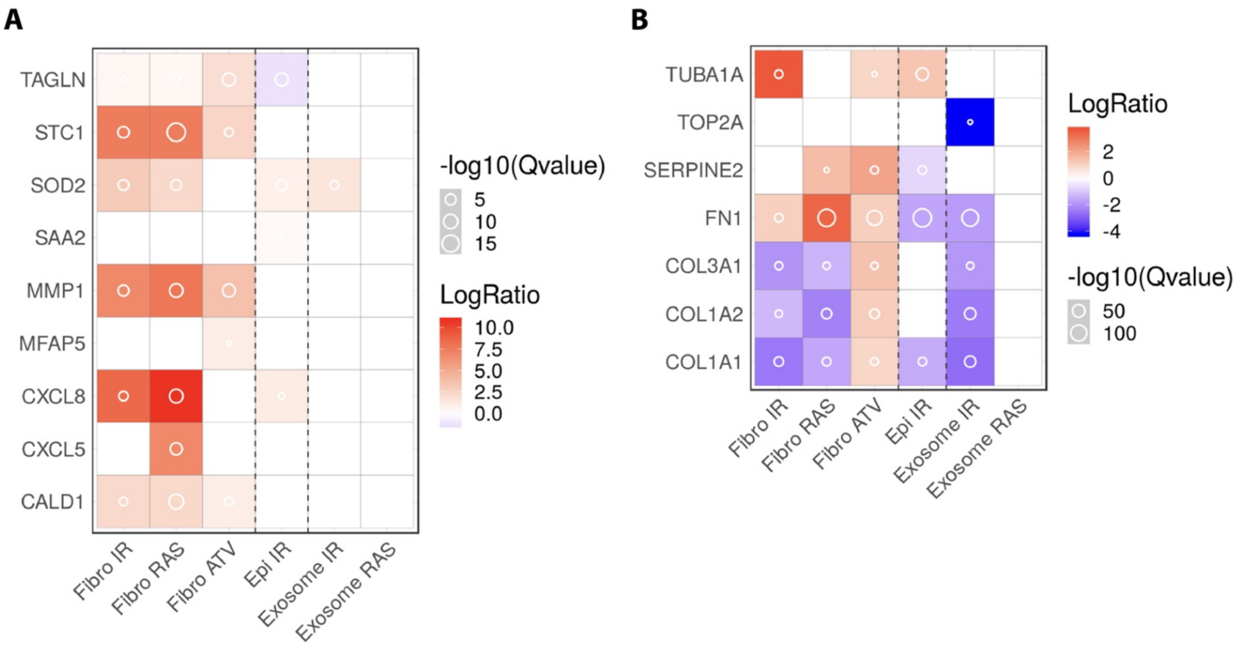


**Supplemental Figure 5**. **Expression of SASP in control and DM1 cells.** Heatmap comparing the genes overexpressed (>1.7 fold) in myoblasts from DM1 patients (**A**) and healthy controls (**B**) to the SASP Atlas (http://www.saspatlas.com/), a proteomic database soluble proteins and exosomes of SASP factors secreted by different cell types: fibroblasts (Fibro) and epithelial cells (Epi) subjected to multiple senescence inducers: genotoxic stress-induced (IR; Irradiation), oncogene-induced (RAS overexpression), treatment-induced (ATV; Atazanivir). Red boxes indicate positive regulators of senescence that are overexpressed by senescent cells of the SASP Atlas. Blue boxes indicate negative regulators of senescence that are overexpressed in non-senescent control cells of the SASP Atlas.

### RNAse negative control

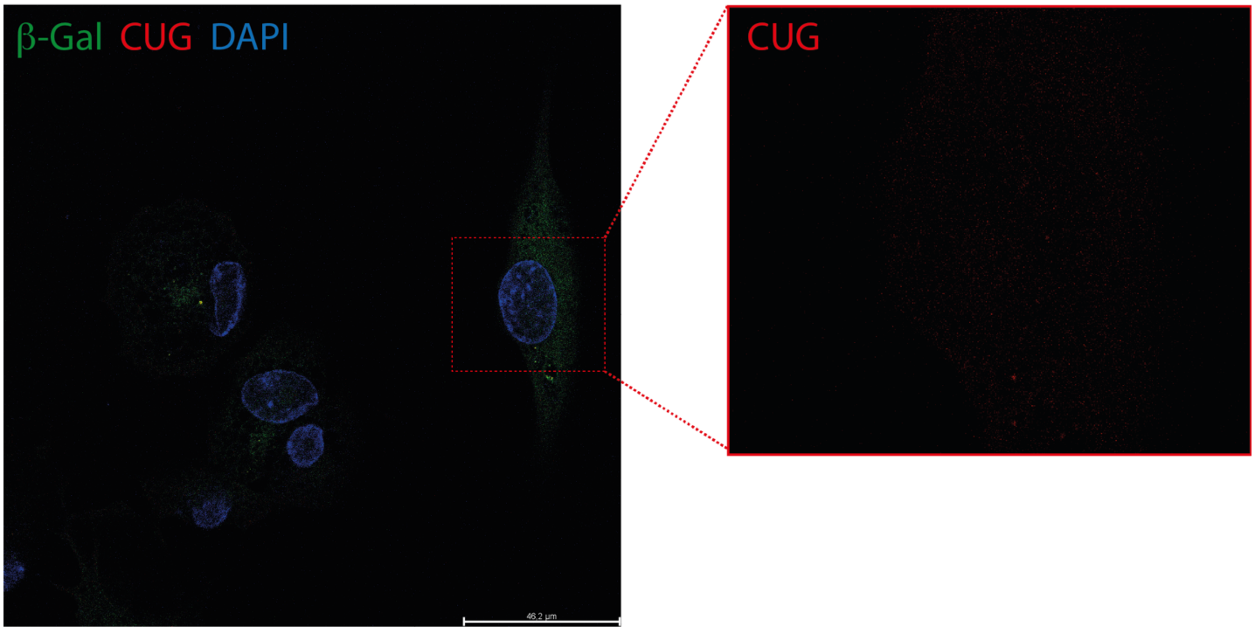


**Supplemental Figure 6: Negative control for detection of RNA foci.** DM1 myoblasts were treated with RNAse as a negative control before performing FISH for detection of CUG repeats. Representative micrograph of immunstaining of SA-β-Gal (green), CUG repeats (red), and DAPI (blue). Intranuclear RNA foci were not detected.

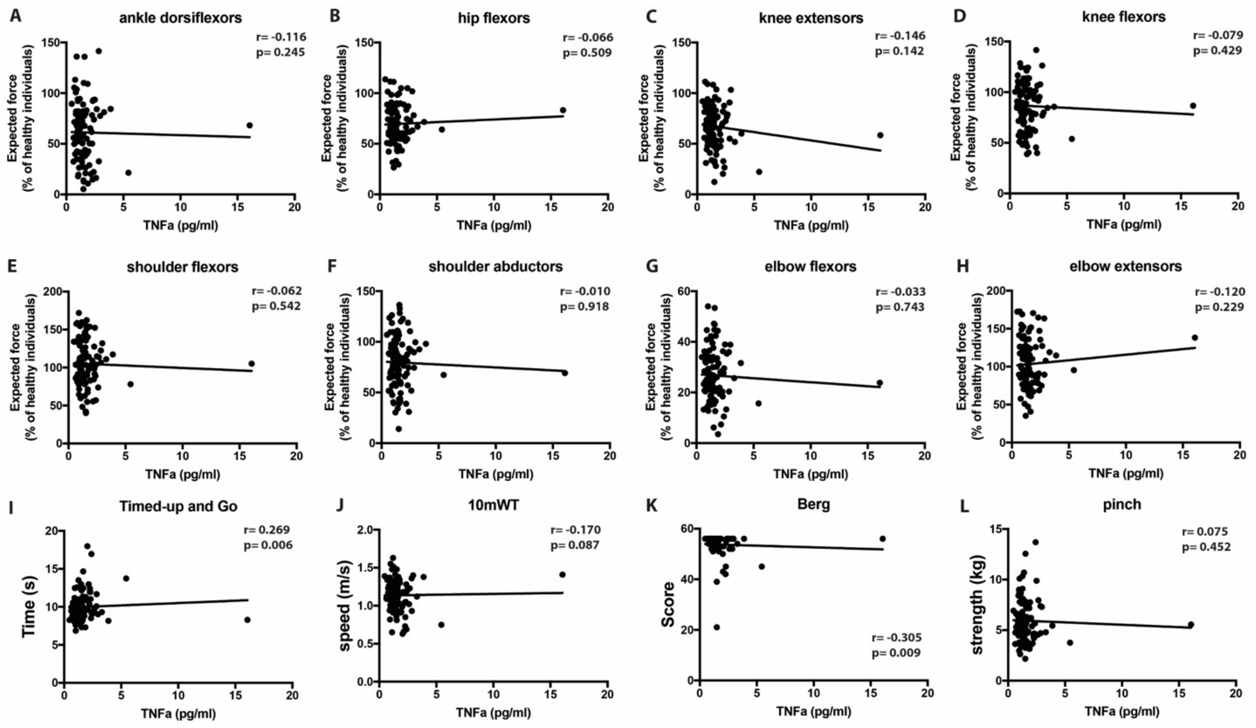


**Supplemental Figure 7. TNFα level is not correlated with muscle strength and functional outcomes in DM1**. Correlation between serum TNFα levels in DM1 patients and the expected strength (relative to normative values) of different muscle groups of **A-D**) the lower limb (ankle dorsiflexors, hip flexors, knee extensors, knee flexors), **E-H**) the upper limb (shoulder flexors, shoulder abductors, elbow flexors, elbow extensors) and **I-L**) different functional capacity tests (Timed-up and Go, 10-meter walk test; 10mWT, Berg Balance scale, pinch test). N=103 patients

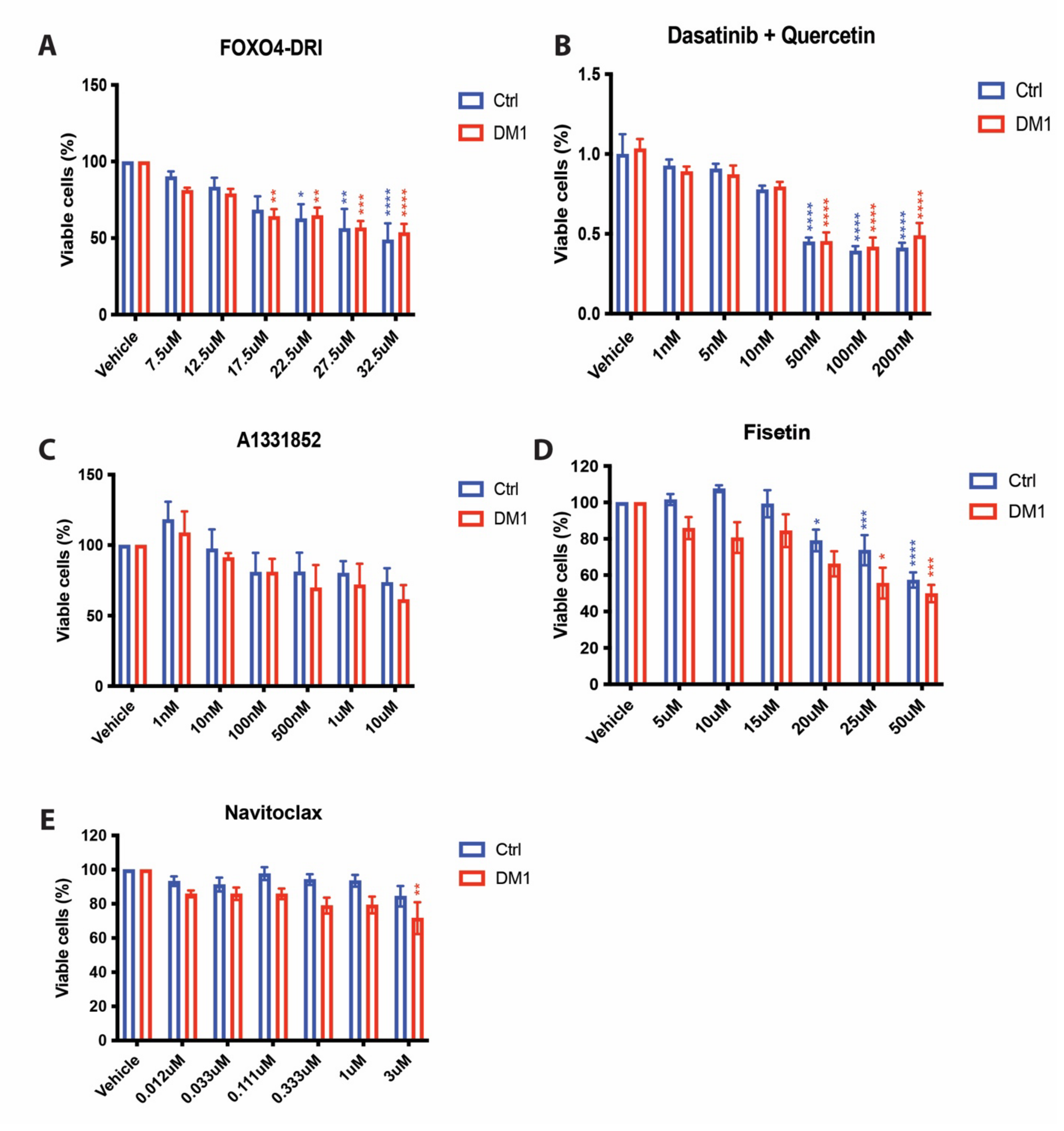


**Supplemental Figure 8**. **Senolytic drug screen**. Cell viability assay of control and DM1 myoblasts treated with **A**) FOXO4-DRI, **B**) dasatinib + quercetin, **C**) A1331852, **D**) fisetin, and **E**) Navitoclax (ABT-263) at different concentrations. Data are expressed as means +/- SEM. N=5. *p<0.05, **p<0.01, ***p<0.001, ****p<0.0001. Red asterisks indicate significant difference compared to DM1 vehicle; blue asterisks indicate significant difference compared to Ctrl vehicle.

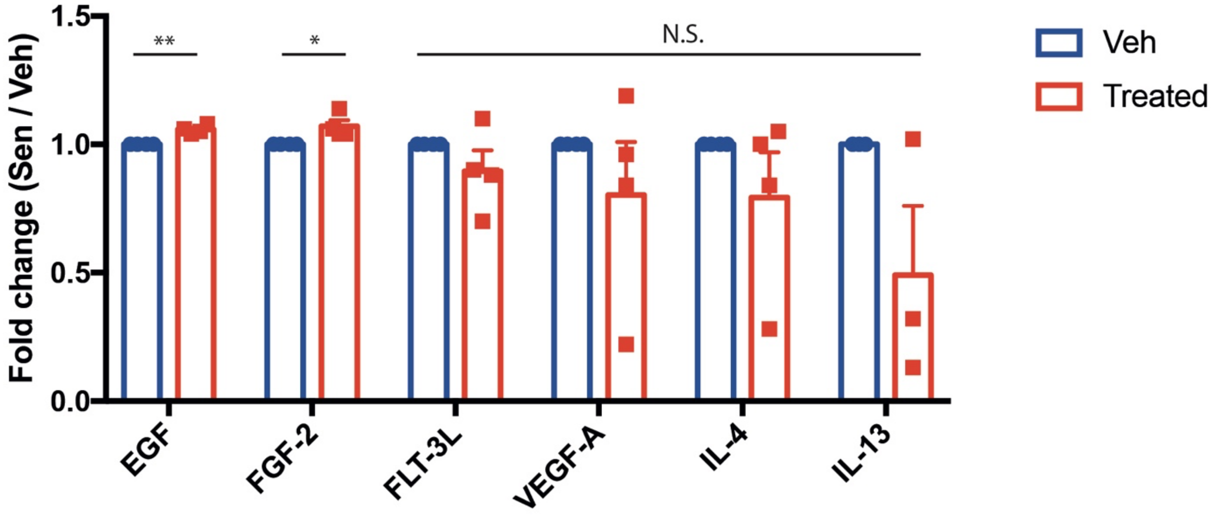


**Supplemental Figure 9. Effect of senolytics on anti-inflammatory cytokines and growth factors.** Multiplex Luminex assay of anti-inflammatory cytokines and growth factors markers found in the supernatant media of DM1 myoblasts treated or not with A1155463. Data are expressed as means +/- SEM. N=4 (except IL-13 treated group: n=3). N.S.: non-significant, *p<0.05, **p<0.01.

| Participant number | Phenotype | CTG (rep) | Sex | Age (y) | scRNAseq |
|---|---|---|---|---|---|
| DM1-452 | Juvenile | 748 | Woman | 58 | No |
| DM1-743 | Adult | 1097 | Woman | 51 | No |
| DM1-1242 | Juvenile | 1200 | Man | 55 | Yes |
| DM1-1594 | Adult | 673 | Man | 58 | No |
| DM1-1621 | Infantile | 675 | Woman | 36 | No |
| DM1-1673 | Juvenile | 554 | Woman | 36 | Yes |
| DM1-1692 | Juvenile | 954 | Man | 35 | Yes |
| DM1-1751 | Juvenile | 1000 | Woman | 30 | No |
| DM1-1770 | Juvenile | 800 | Woman | 32 | No |
| DM1-1948 | Juvenile | 601 | Woman | 30 | No |
| **Mean (SD)** | | **830 (220)** | | **42 (12)** | |
| CTR-4 | N/A | N/A | Man | 33 | Yes |
| CTR-5 | N/A | N/A | Woman | 31 | Yes |
| CTR-7 | N/A | N/A | Woman | 44 | No |
| CTR-9 | N/A | N/A | Woman | 34 | No |
| CTR-10 | N/A | N/A | Man | 45 | No |
| CTR-11 | N/A | N/A | Man | 32 | No |
| CTR-193 | N/A | N/A | Man | 50 | Yes |
| **Mean (SD)** | | **N/A** | | **38.5 (7.5)** | |

**Supplemental table 1.** Clinical characteristics of control and DM1 individuals that provided muscle biopsies. Abbreviations: CTG: cytosine-thymine-guanine; CTR: control subjects; DM1: myotonic dystrophy type 1 participants; rep: repetitions; y: years old; scRNAseq: single-cell RNA sequencing; SD: standard deviation.

| Gene | p_val | avg_log2FC | Control | Patient | p_val_adj |
|---|---|---|---|---|---|
| CXCL5 | 1,4695E-156 | -3,533360522 | 0,518 | 0,839 | 3,678E-152 |
| CXCL8 | 0 | -3,434631561 | 0,233 | 0,848 | 0 |
| CXCL1 | 4,8847E-231 | -2,735642852 | 0,217 | 0,734 | 1,2226E-226 |
| CXCL3 | 1,5858E-174 | -2,597177145 | 0,398 | 0,75 | 3,9692E-170 |
| SAA1 | 2,404E-204 | -2,553815702 | 0,227 | 0,769 | 6,0169E-200 |
| CSF3 | 2,3793E-221 | -2,140614665 | 0,271 | 0,731 | 5,9552E-217 |
| TNNT2 | 3,15582E-46 | -1,998741026 | 0,452 | 0,628 | 7,89869E-42 |
| MMP1 | 2,1901E-268 | -1,953256447 | 0,007 | 0,573 | 5,4816E-264 |
| BEX1 | 7,8406E-193 | -1,667440751 | 0,194 | 0,653 | 1,9624E-188 |
| CXCL2 | 3,2315E-148 | -1,63806753 | 0,215 | 0,603 | 8,088E-144 |
| IL1B | 6,1665E-195 | -1,617018651 | 0,037 | 0,494 | 1,5434E-190 |
| IER3 | 3,01276E-68 | -1,485948408 | 0,77 | 0,763 | 7,54063E-64 |
| TNNI1 | 3,79496E-81 | -1,336065452 | 0,2 | 0,485 | 9,4984E-77 |
| SOD2 | 2,4927E-144 | -1,324468749 | 0,915 | 0,898 | 6,2389E-140 |
| CCND2 | 2,1855E-151 | -1,29106519 | 0,775 | 0,963 | 5,47E-147 |
| SCG5 | 7,0983E-128 | -1,245228316 | 0,515 | 0,744 | 1,7766E-123 |
| MFAP5 | 7,51429E-40 | -1,180545761 | 0,503 | 0,637 | 1,88075E-35 |
| SAA2 | 1,2103E-120 | -1,16908051 | 0,022 | 0,336 | 3,0293E-116 |
| PPME1 | 1,82726E-43 | -1,085048014 | 0,891 | 0,86 | 4,57344E-39 |
| STC1 | 1,95303E-69 | -1,073524299 | 0,381 | 0,595 | 4,88823E-65 |
| TAGLN | 3,41331E-07 | -1,021174569 | 0,949 | 0,953 | 0,008543165 |
| CXCL6 | 1,01459E-91 | -1,020026414 | 0,147 | 0,449 | 2,53941E-87 |
| CCL20 | 7,68751E-97 | -0,991594506 | 0,1 | 0,401 | 1,92411E-92 |
| TNFRSF11B | 8,61092E-34 | -0,948772693 | 0,36 | 0,489 | 2,15523E-29 |
| IL32 | 1,6023E-134 | -0,898840837 | 0,715 | 0,883 | 4,0103E-130 |
| TNNC1 | 1,05146E-79 | -0,890537801 | 0,135 | 0,409 | 2,6317E-75 |
| CALD1 | 9,37747E-22 | -0,878615356 | 0,991 | 0,989 | 2,34709E-17 |
| DDIT3 | 7,67822E-13 | -0,848772749 | 0,659 | 0,693 | 1,92178E-08 |
| INHBA | 3,8931E-19 | -0,845173081 | 0,848 | 0,765 | 9,74403E-15 |
| TNNT1 | 7,78567E-67 | -0,824727422 | 0,158 | 0,406 | 1,94867E-62 |
| MAP1LC3B | 1,7437E-124 | -0,823482564 | 0,975 | 0,971 | 4,3644E-120 |
| ATF3 | 1,33339E-09 | -0,800518342 | 0,559 | 0,6 | 3,33735E-05 |
| ATP6V0C | 6,6265E-126 | -0,761653847 | 0,981 | 0,975 | 1,6585E-121 |
| SLC3A2 | 3,47479E-17 | -0,76069188 | 0,929 | 0,845 | 8,69705E-13 |
| TNFAIP3 | 1,1031E-115 | -0,75987896 | 0,248 | 0,57 | 2,7611E-111 |
| ATP6V0E1 | 2,76705E-68 | -0,739431645 | 0,989 | 0,981 | 6,92565E-64 |
| PPIF | 3,29775E-45 | -0,72336488 | 0,781 | 0,745 | 8,25394E-41 |
| MYL9 | 6,46598E-32 | -0,704676333 | 0,961 | 0,958 | 1,61837E-27 |
| CYTOR | 4,51065E-42 | -0,691782002 | 0,926 | 0,876 | 1,12897E-37 |
| MYL6 | 1,3344E-153 | -0,684196241 | 0,999 | 1 | 3,3399E-149 |
| PPP1R15A | 6,55762E-33 | -0,663759276 | 0,893 | 0,776 | 1,64131E-28 |
| SRGN | 6,54502E-68 | -0,659404241 | 0,949 | 0,964 | 1,63815E-63 |
| PRDX6 | 9,4434E-78 | -0,639848155 | 0,988 | 0,991 | 2,36359E-73 |
| SQSTM1 | 3,87158E-28 | -0,620213589 | 0,963 | 0,905 | 9,69018E-24 |
| LY6K | 5,9992E-63 | -0,616194356 | 0,348 | 0,577 | 1,50154E-58 |
| UBE2B | 2,87347E-52 | -0,611664612 | 0,924 | 0,862 | 7,19202E-48 |
| ATP13A3 | 1,77218E-58 | -0,603109331 | 0,835 | 0,754 | 4,4356E-54 |
| NPC2 | 5,44733E-44 | -0,594825747 | 0,941 | 0,908 | 1,36341E-39 |
| MMP3 | 3,3373E-89 | -0,589973278 | 0,052 | 0,327 | 8,35292E-85 |
| CYCS | 2,5076E-101 | -0,589127467 | 0,976 | 0,97 | 6,27635E-97 |
| GREM1 | 5,30977E-21 | -0,589001958 | 0,934 | 0,902 | 1,32898E-16 |
| ATP6V1G1 | 4,68887E-48 | -0,585223822 | 0,979 | 0,976 | 1,17358E-43 |
| MGLL | 3,8528E-106 | -0,579317715 | 0,855 | 0,896 | 9,6433E-102 |
| SPHK1 | 4,1366E-54 | -0,57564691 | 0,572 | 0,68 | 1,03535E-49 |
| ATP6V0B | 2,31401E-85 | -0,568316311 | 0,963 | 0,926 | 5,79172E-81 |
| IGFBP5 | 1,99872E-65 | -0,565761587 | 0,02 | 0,212 | 5,0026E-61 |
| UCHL1 | 1,09615E-21 | -0,565735338 | 0,672 | 0,716 | 2,74356E-17 |
| UBC | 3,93416E-14 | -0,559499748 | 0,998 | 0,992 | 9,84681E-10 |
| B2M | 5,52268E-19 | -0,548383789 | 0,997 | 0,995 | 1,38227E-14 |
| POU2F2 | 2,6795E-141 | -0,544553003 | 0,066 | 0,435 | 6,7066E-137 |
| GABARAPL2 | 1,09524E-94 | -0,539567897 | 0,975 | 0,958 | 2,74127E-90 |
| AKR1B1 | 6,25246E-75 | -0,532568034 | 0,944 | 0,89 | 1,56493E-70 |
| HLA-B | 1,20013E-39 | -0,529068056 | 0,883 | 0,871 | 3,00381E-35 |
| LUM | 8,83668E-55 | -0,526894707 | 0,388 | 0,627 | 2,21173E-50 |
| PMAIP1 | 1,0907E-25 | -0,522359599 | 0,535 | 0,592 | 2,7299E-21 |
| TPM2 | 4,59934E-31 | -0,521471652 | 0,997 | 0,992 | 1,15117E-26 |
| RNASEK | 1,92237E-74 | -0,519321265 | 0,993 | 0,991 | 4,8115E-70 |
| FGF2 | 4,07408E-36 | -0,51093067 | 0,765 | 0,768 | 1,0197E-31 |
| MOK | 2,54199E-80 | -0,500618574 | 0,61 | 0,71 | 6,36234E-76 |
| MYL12B | 9,15728E-54 | -0,49976052 | 0,997 | 0,994 | 2,29198E-49 |
| SRP14 | 3,75731E-38 | -0,495805338 | 0,999 | 0,997 | 9,40418E-34 |
| BGN | 4,9499E-71 | -0,495606748 | 0,147 | 0,393 | 1,23891E-66 |
| PAPPA | 5,90915E-45 | -0,495533104 | 0,359 | 0,55 | 1,479E-40 |
| NDUFV3 | 8,0358E-153 | -0,49082777 | 0,947 | 0,915 | 2,0113E-148 |
| POLR2L | 1,91433E-66 | -0,489888808 | 0,997 | 0,996 | 4,79138E-62 |
| SAP18 | 1,4066E-90 | -0,487782492 | 0,988 | 0,977 | 3,52057E-86 |
| MYL12A | 1,44447E-53 | -0,486415143 | 0,996 | 0,989 | 3,61535E-49 |
| OPTN | 5,9598E-109 | -0,485783883 | 0,906 | 0,919 | 1,4917E-104 |
| NFKBIZ | 1,01582E-14 | -0,484082987 | 0,75 | 0,668 | 2,54249E-10 |
| ADM | 8,55806E-27 | -0,481322094 | 0,707 | 0,707 | 2,142E-22 |
| ITM2B | 3,83603E-20 | -0,472902654 | 0,965 | 0,933 | 9,60121E-16 |
| CLDN11 | 4,10341E-30 | -0,472028658 | 0,331 | 0,473 | 1,02704E-25 |
| STEAP1 | 1,01476E-06 | -0,460822799 | 0,626 | 0,546 | 0,025398461 |
| RHOC | 4,14247E-89 | -0,451759912 | 0,983 | 0,969 | 1,03682E-84 |
| TNFSF4 | 1,78278E-07 | -0,447070457 | 0,368 | 0,399 | 0,004462118 |
| SKP1 | 5,86099E-46 | -0,443056754 | 0,996 | 0,995 | 1,46695E-41 |
| S100A4 | 1,74995E-06 | -0,44275265 | 0,981 | 0,96 | 0,043799457 |
| ASAH1 | 2,74524E-15 | -0,441238244 | 0,913 | 0,861 | 6,87106E-11 |
| TSC22D1 | 9,45305E-19 | -0,439359459 | 0,822 | 0,73 | 2,366E-14 |
| MAP3K7CL | 1,34077E-14 | -0,435529736 | 0,687 | 0,685 | 3,35581E-10 |
| NOP10 | 1,80572E-80 | -0,434786573 | 0,993 | 0,981 | 4,51954E-76 |
| CD99 | 3,13681E-42 | -0,431135591 | 0,978 | 0,96 | 7,85113E-38 |
| RPS26 | 4,3724E-85 | -0,43022251 | 0,998 | 0,999 | 1,09437E-80 |
| ELOC | 5,0678E-107 | -0,430000499 | 0,984 | 0,97 | 1,2684E-102 |
| HHIP | 1,51762E-06 | -0,427457434 | 0,352 | 0,368 | 0,037984413 |
| IL6 | 1,65279E-46 | -0,427432856 | 0,133 | 0,328 | 4,13676E-42 |
| TBCA | 5,5768E-105 | -0,423616427 | 0,989 | 0,982 | 1,3958E-100 |
| TNFRSF10D | 9,33726E-15 | -0,416362587 | 0,802 | 0,754 | 2,33702E-10 |
| EDN1 | 1,63962E-08 | -0,411774251 | 0,447 | 0,51 | 0,000410381 |
| KRTAP2-3 | 2,21118E-25 | -0,410126049 | 0,201 | 0,354 | 5,53436E-21 |
| UBB | 7,79194E-25 | -0,408776885 | 0,997 | 0,992 | 1,95024E-20 |
| ANGPTL4 | 2,24005E-07 | -0,404961521 | 0,709 | 0,679 | 0,005606623 |
| AL118516.1 | 2,17672E-10 | -0,404915409 | 0,49 | 0,574 | 5,44812E-06 |
| PSAP | 5,84039E-18 | -0,39850935 | 0,952 | 0,881 | 1,46179E-13 |

| | | | | | |
|---|---|---|---|---|---|
| **MRPL33** | 5,88037E-85 | -0,397057196 | 0,982 | 0,967 | 1,4718E-80 |
| **MOCS2** | 1,74966E-54 | -0,393310234 | 0,92 | 0,905 | 4,37922E-50 |
| **SOD1** | 1,60874E-61 | -0,393289746 | 0,995 | 0,989 | 4,02651E-57 |
| **ARF4** | 4,83127E-88 | -0,392386603 | 0,984 | 0,973 | 1,20922E-83 |
| **ANAPC13** | 2,68581E-71 | -0,391014548 | 0,949 | 0,917 | 6,72231E-67 |
| **GPNMB** | 1,47139E-18 | -0,390466072 | 0,43 | 0,534 | 3,68273E-14 |
| **CALU** | 7,11052E-14 | -0,389368554 | 0,983 | 0,975 | 1,77969E-09 |
| **FKBP1A** | 4,52541E-39 | -0,388808948 | 0,986 | 0,966 | 1,13267E-34 |
| **SERINC2** | 1,96579E-52 | -0,388595634 | 0,804 | 0,817 | 4,92018E-48 |
| **SDCBP** | 1,40304E-34 | -0,386003893 | 0,963 | 0,93 | 3,51168E-30 |
| **PADI2** | 3,5515E-46 | -0,384337955 | 0,009 | 0,141 | 8,88904E-42 |
| **CHIC2** | 7,84304E-18 | -0,380685743 | 0,827 | 0,745 | 1,96304E-13 |
| **FTH1** | 2,65238E-41 | -0,378296074 | 1 | 1 | 6,63865E-37 |
| **SERPINE1** | 2,41647E-09 | -0,377790347 | 0,994 | 0,989 | 6,04819E-05 |
| **COX6C** | 8,81361E-52 | -0,372987163 | 0,99 | 0,986 | 2,20596E-47 |
| **UBL5** | 1,03978E-46 | -0,371748921 | 0,994 | 0,994 | 2,60246E-42 |
| **S100A11** | 1,09229E-31 | -0,367502165 | 1 | 0,998 | 2,73388E-27 |
| **COX7B** | 1,26696E-61 | -0,367228314 | 0,993 | 0,98 | 3,17107E-57 |
| **CLTB** | 2,01308E-43 | -0,366015529 | 0,923 | 0,893 | 5,03854E-39 |
| **OGFRL1** | 3,01354E-29 | -0,364893658 | 0,779 | 0,784 | 7,54259E-25 |
| **ATF4** | 3,42373E-07 | -0,363155408 | 0,963 | 0,881 | 0,008569243 |
| **SERINC1** | 3,02467E-09 | -0,362484267 | 0,953 | 0,922 | 7,57045E-05 |
| **CDA** | 3,11403E-92 | -0,357598336 | 0,162 | 0,455 | 7,79412E-88 |
| **CSTB** | 3,07889E-34 | -0,355901697 | 0,988 | 0,967 | 7,70615E-30 |
| **MGST1** | 2,13273E-25 | -0,354215785 | 0,879 | 0,912 | 5,33802E-21 |
| **PHLDA2** | 1,80161E-08 | -0,351381549 | 0,864 | 0,804 | 0,000450924 |
| **NDUFB3** | 2,87938E-55 | -0,350248428 | 0,989 | 0,975 | 7,2068E-51 |
| **LINC02802** | 2,8667E-45 | -0,350108564 | 0,961 | 0,905 | 7,17506E-41 |
| **SEC11C** | 1,40858E-69 | -0,348394979 | 0,852 | 0,842 | 3,52554E-65 |
| **MPC2** | 9,19942E-43 | -0,346391395 | 0,944 | 0,887 | 2,30252E-38 |
| **BST2** | 6,4071E-87 | -0,346352157 | 0,098 | 0,376 | 1,60363E-82 |
| **ODC1** | 2,37539E-13 | -0,345235224 | 0,904 | 0,814 | 5,94535E-09 |
| **DYNLRB1** | 1,07991E-57 | -0,34253708 | 0,983 | 0,975 | 2,7029E-53 |
| **TXNIP** | 3,22837E-15 | -0,342363273 | 0,863 | 0,692 | 8,08028E-11 |
| **EIF5** | 5,19233E-08 | -0,340761617 | 0,933 | 0,842 | 0,001299588 |
| **NDUFA2** | 3,40721E-40 | -0,339212766 | 0,981 | 0,962 | 8,5279E-36 |
| **RNF114** | 6,88131E-11 | -0,338725724 | 0,761 | 0,724 | 1,72232E-06 |
| **MAP2K3** | 2,09321E-40 | -0,334487072 | 0,823 | 0,757 | 5,23911E-36 |
| **SELENOK** | 5,50735E-55 | -0,331410846 | 0,967 | 0,942 | 1,37844E-50 |
| **PPDPF** | 4,73078E-07 | -0,326656927 | 0,994 | 0,985 | 0,011840669 |
| **NQO1** | 1,04771E-19 | -0,325676153 | 0,98 | 0,957 | 2,6223E-15 |
| **SAT1** | 2,97172E-11 | -0,325517072 | 0,975 | 0,931 | 7,43791E-07 |
| **TXNRD1** | 3,66824E-13 | -0,324728903 | 0,948 | 0,864 | 9,18124E-09 |
| **BRI3** | 4,3275E-17 | -0,32393567 | 0,957 | 0,921 | 1,08313E-12 |
| **MCOLN1** | 5,2793E-07 | -0,323726284 | 0,664 | 0,671 | 0,01321355 |
| **REEP5** | 9,6968E-59 | -0,32358687 | 0,973 | 0,947 | 2,42701E-54 |
| **NGFR** | 5,74554E-12 | -0,321596502 | 0,169 | 0,26 | 1,43805E-07 |
| **ATP6V0D1** | 7,9495E-22 | -0,321422401 | 0,926 | 0,849 | 1,98968E-17 |
| **ATP6V1D** | 7,21094E-34 | -0,320395979 | 0,975 | 0,933 | 1,80483E-29 |
| **POMP** | 7,14637E-73 | -0,320361557 | 0,991 | 0,987 | 1,78867E-68 |
| **NDUFA1** | 5,37207E-41 | -0,320014135 | 0,991 | 0,987 | 1,34458E-36 |
| **SH3BGRL3** | 2,239E-14 | -0,318697782 | 0,998 | 0,994 | 5,604E-10 |
| **ATP6V1C1** | 1,27591E-09 | -0,31749187 | 0,85 | 0,766 | 3,19348E-05 |
| **YPEL5** | 1,30208E-21 | -0,315015167 | 0,848 | 0,803 | 3,25898E-17 |
| **CHST2** | 2,29582E-09 | -0,314254083 | 0,591 | 0,617 | 5,7462E-05 |
| **KRT7** | 4,85307E-15 | -0,31315887 | 0,632 | 0,685 | 1,21468E-10 |
| **DSTN** | 6,0497E-36 | -0,312899298 | 0,988 | 0,985 | 1,51418E-31 |
| **MIR4435-2HG** | 1,58075E-21 | -0,31254906 | 0,886 | 0,796 | 3,95646E-17 |
| **NEK7** | 6,65204E-09 | -0,312457863 | 0,815 | 0,796 | 0,000166494 |
| **TNFAIP2** | 8,4347E-140 | -0,311320566 | 0,189 | 0,597 | 2,1111E-135 |
| **PNP** | 4,68119E-13 | -0,309886942 | 0,794 | 0,718 | 1,17165E-08 |
| **TAF13** | 1,01217E-70 | -0,309386196 | 0,871 | 0,859 | 2,53335E-66 |
| **COL6A3** | 2,63824E-07 | -0,309069935 | 0,88 | 0,714 | 0,00660326 |
| **ATP5MD** | 6,12284E-45 | -0,308216833 | 0,996 | 0,988 | 1,53249E-40 |
| **C15orf48** | 8,82967E-93 | -0,307928619 | 0,019 | 0,269 | 2,20998E-88 |
| **GABARAP** | 6,05899E-17 | -0,307734214 | 0,988 | 0,984 | 1,5165E-12 |
| **IFRD1** | 2,4984E-12 | -0,307401647 | 0,764 | 0,711 | 6,25325E-08 |
| **ATP5MG** | 8,27279E-52 | -0,306707194 | 0,995 | 0,987 | 2,0706E-47 |
| **CHMP5** | 1,6091E-32 | -0,305787212 | 0,979 | 0,961 | 4,02741E-28 |
| **ESF1** | 1,19706E-08 | -0,304976519 | 0,92 | 0,818 | 0,000299611 |
| **COX6A1** | 1,69671E-28 | -0,303443905 | 0,994 | 0,986 | 4,2467E-24 |
| **POSTN** | 1,58729E-11 | -0,303007896 | 0,307 | 0,402 | 3,97283E-07 |
| **COX7A2** | 2,02852E-52 | -0,302355111 | 0,993 | 0,989 | 5,07718E-48 |
| **IL7R** | 8,14682E-09 | -0,3022058 | 0,555 | 0,592 | 0,000203907 |
| **RIT1** | 5,79351E-13 | -0,301136252 | 0,794 | 0,741 | 1,45006E-08 |
| **VAT1** | 3,68591E-38 | -0,299860792 | 0,911 | 0,885 | 9,22545E-34 |
| **COX7A2L** | 3,00798E-61 | -0,299108242 | 0,98 | 0,959 | 7,52867E-57 |
| **ATP6V1F** | 1,92854E-36 | -0,298217567 | 0,978 | 0,962 | 4,82693E-32 |
| **NDUFS5** | 2,96613E-43 | -0,296466962 | 0,991 | 0,987 | 7,42393E-39 |
| **COX6B1** | 7,94343E-40 | -0,296241399 | 0,993 | 0,981 | 1,98816E-35 |
| **SEC61G** | 1,46703E-43 | -0,295328104 | 0,994 | 0,987 | 3,67183E-39 |
| **CES1** | 5,20586E-53 | -0,294896545 | 0,022 | 0,183 | 1,30297E-48 |
| **IFI27** | 1,06648E-26 | -0,29339179 | 0,168 | 0,323 | 2,66929E-22 |
| **UQCR11** | 4,18614E-31 | -0,29237808 | 0,992 | 0,987 | 1,04775E-26 |
| **CKB** | 1,33862E-37 | -0,292077212 | 0,24 | 0,43 | 3,35044E-33 |
| **SORBS2** | 2,79962E-39 | -0,291454053 | 0,342 | 0,514 | 7,00717E-35 |
| **UPP1** | 1,31721E-17 | -0,290599279 | 0,641 | 0,693 | 3,29685E-13 |
| **HIGD1A** | 1,00766E-50 | -0,289141907 | 0,965 | 0,93 | 2,52208E-46 |
| **ATP5MF** | 3,55189E-35 | -0,288815263 | 0,991 | 0,988 | 8,89003E-31 |
| **NDUFA4** | 1,36993E-17 | -0,288707759 | 0,997 | 0,989 | 3,42879E-13 |
| **RHEB** | 5,24748E-11 | -0,288707697 | 0,946 | 0,884 | 1,31339E-06 |
| **C3** | 1,6074E-159 | -0,286532432 | 0,029 | 0,42 | 4,0232E-155 |
| **TAF9** | 3,19701E-30 | -0,286300794 | 0,954 | 0,89 | 8,00181E-26 |
| **GTF2A2** | 9,74402E-38 | -0,285494445 | 0,973 | 0,942 | 2,43883E-33 |
| **TIMP3** | 6,90166E-11 | -0,285104109 | 0,75 | 0,596 | 1,72742E-06 |
| **NEDD8** | 5,00745E-32 | -0,284763291 | 0,991 | 0,978 | 1,25332E-27 |
| **RAB9A** | 7,4134E-15 | -0,28456449 | 0,754 | 0,711 | 1,8555E-10 |
| **ATOX1** | 7,28403E-36 | -0,282523414 | 0,983 | 0,966 | 1,82312E-31 |
| **HSPB3** | 1,20375E-12 | -0,282394017 | 0,29 | 0,374 | 3,01288E-08 |
| **TRAPPC4** | 6,75652E-45 | -0,282069266 | 0,911 | 0,817 | 1,69109E-40 |
| **DKK3** | 9,01441E-12 | -0,281887794 | 0,653 | 0,665 | 2,25622E-07 |
| **PFDN2** | 5,64694E-34 | -0,281740465 | 0,973 | 0,932 | 1,41337E-29 |
| **JAK1** | 4,6901E-39 | -0,281453525 | 0,929 | 0,884 | 1,17388E-34 |
| **FAM177A1** | 2,95046E-59 | -0,280840597 | 0,96 | 0,928 | 7,3847E-55 |
| **RALA** | 5,26988E-14 | -0,279536702 | 0,933 | 0,84 | 1,319E-09 |

| | | | | | |
|---|---|---|---|---|---|
| UQCRQ | 8,37603E-24 | -0,279412099 | 0,99 | 0,982 | 2,09644E-19 |
| CHCHD2 | 2,85881E-61 | -0,279127667 | 0,996 | 0,991 | 7,15533E-57 |
| AC015912.3 | 8,38144E-51 | -0,278492803 | 0,191 | 0,419 | 2,09779E-46 |
| PRDX1 | 1,10607E-24 | -0,277557099 | 0,998 | 0,996 | 2,76838E-20 |
| BRK1 | 1,18077E-43 | -0,277432012 | 0,979 | 0,977 | 2,95534E-39 |
| CUTA | 2,11153E-15 | -0,277142504 | 0,986 | 0,966 | 5,28494E-11 |
| MXRA7 | 2,05615E-13 | -0,275612988 | 0,953 | 0,933 | 5,14634E-09 |
| ROMO1 | 1,88436E-18 | -0,274824131 | 0,989 | 0,973 | 4,71636E-14 |
| PHPT1 | 6,22672E-10 | -0,274076267 | 0,99 | 0,975 | 1,55849E-05 |
| IL13RA2 | 1,78071E-27 | -0,27367375 | 0,134 | 0,274 | 4,45694E-23 |
| GRPEL1 | 4,06814E-43 | -0,273553999 | 0,933 | 0,852 | 1,01821E-38 |
| ATP5IF1 | 4,89422E-29 | -0,271270295 | 0,99 | 0,985 | 1,22498E-24 |
| DNAJB9 | 6,87563E-08 | -0,27101151 | 0,702 | 0,687 | 0,0017209 |
| ZNF57 | 3,07728E-09 | -0,270739807 | 0,102 | 0,17 | 7,70212E-05 |
| PRNP | 3,88268E-11 | -0,269633669 | 0,861 | 0,81 | 9,71796E-07 |
| SBDS | 8,02066E-40 | -0,268018379 | 0,976 | 0,955 | 2,00749E-35 |
| IGFBP2 | 5,56715E-24 | -0,266467467 | 0,127 | 0,265 | 1,3934E-19 |
| PTS | 4,2851E-42 | -0,266432337 | 0,913 | 0,857 | 1,07252E-37 |
| CAVIN4 | 6,15243E-13 | -0,26621878 | 0,237 | 0,329 | 1,53989E-08 |
| CCT8 | 6,87353E-33 | -0,265534285 | 0,983 | 0,96 | 1,72038E-28 |
| MAFF | 2,13244E-38 | -0,265312432 | 0,596 | 0,671 | 5,33729E-34 |
| EPB41L3 | 4,2525E-133 | -0,264693356 | 0,009 | 0,334 | 1,0644E-128 |
| NDUFB5 | 9,24091E-47 | -0,264525379 | 0,964 | 0,933 | 2,31291E-42 |
| CTSL | 1,8105E-18 | -0,264350711 | 0,952 | 0,86 | 4,53151E-14 |
| ADAMTS12 | 1,20064E-70 | -0,264323793 | 0,402 | 0,623 | 3,00509E-66 |
| QSOX1 | 9,62871E-09 | -0,263979059 | 0,884 | 0,768 | 0,000240997 |
| HBEGF | 2,26764E-71 | -0,262991337 | 0,28 | 0,538 | 5,67568E-67 |
| SEC22B | 1,472E-43 | -0,260993173 | 0,96 | 0,908 | 3,68428E-39 |
| NAMPT | 3,69581E-08 | -0,260578156 | 0,869 | 0,759 | 0,000925025 |
| EIF1AY | 5,9955E-23 | -0,260147203 | 0,478 | 0,586 | 1,50061E-18 |
| CSNK2B | 3,6548E-54 | -0,259287544 | 0,982 | 0,943 | 9,1476E-50 |
| MAP1LC3A | 3,43081E-32 | -0,257506327 | 0,76 | 0,777 | 8,58697E-28 |
| UFM1 | 9,32863E-50 | -0,257372181 | 0,965 | 0,929 | 2,33486E-45 |
| VAMP5 | 2,23449E-08 | -0,256327345 | 0,973 | 0,926 | 0,000559271 |
| SUB1 | 1,52566E-48 | -0,256084427 | 0,989 | 0,978 | 3,81859E-44 |
| RNF181 | 2,9105E-10 | -0,25578908 | 0,981 | 0,957 | 7,2847E-06 |
| PDLIM1 | 7,03839E-79 | -0,255447459 | 0,372 | 0,645 | 1,76164E-74 |
| TIMM13 | 2,67131E-51 | -0,254148723 | 0,975 | 0,949 | 6,68603E-47 |
| NDUFC1 | 1,91199E-36 | -0,25373754 | 0,981 | 0,962 | 4,78552E-32 |
| COA6 | 1,04742E-10 | -0,250339025 | 0,933 | 0,83 | 2,6216E-06 |
| NRG1 | 2,77325E-16 | -0,250319624 | 0,543 | 0,621 | 6,94116E-12 |
| PTPN14 | 1,70799E-64 | 0,250578943 | 0,889 | 0,718 | 4,27492E-60 |
| PLXNA2 | 1,13494E-65 | 0,250688551 | 0,798 | 0,598 | 2,84065E-61 |
| JPT1 | 1,93879E-16 | 0,251140036 | 0,97 | 0,894 | 4,85261E-12 |
| HP1BP3 | 1,08797E-47 | 0,25114794 | 0,944 | 0,833 | 2,72309E-43 |
| LASP1 | 2,47271E-62 | 0,251399482 | 0,91 | 0,747 | 6,18894E-58 |
| TMSB15A | 1,4639E-103 | 0,251678757 | 0,434 | 0,119 | 3,664E-99 |
| ARL2 | 7,93648E-86 | 0,252183967 | 0,928 | 0,755 | 1,98642E-81 |
| LAMA2 | 4,85027E-97 | 0,253049219 | 0,793 | 0,523 | 1,21397E-92 |
| PF4 | 1,57452E-80 | 0,253146925 | 0,677 | 0,378 | 3,94086E-76 |
| GLUL | 2,84496E-73 | 0,253695996 | 0,869 | 0,692 | 7,12064E-69 |
| CDK1 | 1,18289E-17 | 0,253750358 | 0,354 | 0,231 | 2,96064E-13 |
| SSRP1 | 3,25947E-45 | 0,253883384 | 0,913 | 0,752 | 8,15812E-41 |
| CAMK2N1 | 2,19033E-54 | 0,254162331 | 0,877 | 0,735 | 5,48218E-50 |
| SEC31A | 8,31492E-47 | 0,254282711 | 0,967 | 0,89 | 2,08114E-42 |
| MT-ND5 | 3,94139E-40 | 0,254379424 | 0,981 | 0,938 | 9,8649E-36 |
| RDX | 1,86485E-37 | 0,254906894 | 0,959 | 0,91 | 4,66753E-33 |
| MYH10 | 7,96516E-89 | 0,255615047 | 0,703 | 0,454 | 1,9936E-84 |
| RAPSN | 6,36222E-67 | 0,255936427 | 0,8 | 0,525 | 1,5924E-62 |
| GTF2I | 1,55268E-65 | 0,256052209 | 0,918 | 0,777 | 3,88621E-61 |
| HIP1 | 1,77983E-84 | 0,256975556 | 0,78 | 0,567 | 4,45475E-80 |
| PDLIM3 | 2,40181E-63 | 0,257534752 | 0,846 | 0,561 | 6,01149E-59 |
| FERMT2 | 6,41507E-40 | 0,258934338 | 0,918 | 0,775 | 1,60563E-35 |
| TACC3 | 6,48173E-21 | 0,259143149 | 0,42 | 0,304 | 1,62231E-16 |
| LMNB1 | 1,80046E-49 | 0,259245483 | 0,404 | 0,202 | 4,50638E-45 |
| HNRNPM | 1,58805E-41 | 0,259616076 | 0,97 | 0,908 | 3,97474E-37 |
| MAP3K5 | 8,4569E-147 | 0,259867344 | 0,742 | 0,397 | 2,1167E-142 |
| PLPP3 | 1,77988E-46 | 0,261835064 | 0,814 | 0,624 | 4,45486E-42 |
| ARHGEF28 | 9,91465E-90 | 0,262471278 | 0,78 | 0,558 | 2,48154E-85 |
| KPNB1 | 9,82809E-33 | 0,262916637 | 0,955 | 0,888 | 2,45987E-28 |
| ITGA10 | 3,02051E-75 | 0,263497752 | 0,723 | 0,48 | 7,56003E-71 |
| FUS | 2,38768E-33 | 0,263556364 | 0,931 | 0,793 | 5,97613E-29 |
| SPDL1 | 3,7352E-38 | 0,263834119 | 0,851 | 0,701 | 9,34882E-34 |
| ATOH8 | 8,5403E-126 | 0,264354104 | 0,775 | 0,463 | 2,1376E-121 |
| MIOS | 7,9003E-113 | 0,264994809 | 0,799 | 0,538 | 1,9774E-108 |
| KLF9 | 2,1698E-78 | 0,265017127 | 0,846 | 0,623 | 5,43078E-74 |
| ASAP2 | 1,31517E-77 | 0,26699169 | 0,866 | 0,693 | 3,29173E-73 |
| GNAI2 | 5,70823E-53 | 0,268980391 | 0,969 | 0,9 | 1,42871E-48 |
| H2AFV | 9,81857E-13 | 0,269738235 | 0,941 | 0,88 | 2,45749E-08 |
| MT-CO3 | 9,19801E-42 | 0,270624234 | 0,987 | 0,954 | 2,30217E-37 |
| DNAJC9 | 1,15482E-35 | 0,271304898 | 0,807 | 0,669 | 2,8904E-31 |
| XPO1 | 1,01641E-48 | 0,271548675 | 0,913 | 0,736 | 2,54397E-44 |
| TRAM1 | 3,05227E-39 | 0,272713959 | 0,954 | 0,848 | 7,63954E-35 |
| HERPUD1 | 3,94366E-59 | 0,274261154 | 0,926 | 0,771 | 9,87059E-55 |
| FSCN1 | 1,27614E-74 | 0,275034301 | 0,929 | 0,79 | 3,19405E-70 |
| PLEC | 3,22821E-48 | 0,275167628 | 0,916 | 0,786 | 8,07988E-44 |
| SPOCK1 | 1,78047E-77 | 0,27628119 | 0,825 | 0,634 | 4,45635E-73 |
| TPM4 | 5,34012E-33 | 0,276390234 | 0,994 | 0,974 | 1,33658E-28 |
| NREP | 2,34904E-49 | 0,276707737 | 0,884 | 0,753 | 5,87941E-45 |
| KNL1 | 3,05171E-31 | 0,27796004 | 0,386 | 0,222 | 7,63812E-27 |
| MARCKSL1 | 6,14758E-78 | 0,277977739 | 0,762 | 0,541 | 1,53868E-73 |
| HSPG2 | 2,07176E-83 | 0,278690333 | 0,867 | 0,657 | 5,18541E-79 |
| MT2A | 1,21687E-18 | 0,279286274 | 1 | 1 | 3,0457E-14 |
| RBMX | 4,84445E-38 | 0,280322985 | 0,943 | 0,841 | 1,21252E-33 |
| CTHRC1 | 7,47354E-84 | 0,280358481 | 0,693 | 0,438 | 1,87055E-79 |
| NCAPD2 | 2,79904E-59 | 0,280570395 | 0,661 | 0,453 | 7,00573E-55 |
| ACTG1 | 1,09787E-31 | 0,280664705 | 0,998 | 0,99 | 2,74786E-27 |
| ACTR2 | 1,30833E-58 | 0,280991255 | 0,973 | 0,926 | 3,27462E-54 |
| CCNB2 | 2,01999E-31 | 0,281074284 | 0,437 | 0,275 | 5,05583E-27 |
| KNOP1 | 1,01299E-70 | 0,28142344 | 0,893 | 0,687 | 2,5354E-66 |
| DHFR | 8,06283E-30 | 0,28178917 | 0,683 | 0,552 | 2,01805E-25 |
| RBMS1 | 3,43646E-45 | 0,282958676 | 0,917 | 0,744 | 8,60111E-41 |
| TMSB4X | 1,82569E-60 | 0,283592818 | 0,999 | 0,997 | 4,56951E-56 |
| APEX1 | 1,21887E-54 | 0,284205562 | 0,959 | 0,883 | 3,05072E-50 |
| CLMP | 5,57484E-50 | 0,284769891 | 0,884 | 0,689 | 1,39533E-45 |

| | | | | | |
|---|---|---|---|---|---|
| **CENPU** | 7,71977E-33 | 0,284770125 | 0,439 | 0,294 | 1,93218E-28 |
| **MCM7** | 1,96727E-28 | 0,28480293 | 0,628 | 0,491 | 4,92387E-24 |
| **NAV1** | 1,87866E-58 | 0,285728173 | 0,882 | 0,695 | 4,70209E-54 |
| **TK1** | 2,08825E-43 | 0,287027916 | 0,488 | 0,277 | 5,22667E-39 |
| **SLC8A1** | 5,4447E-96 | 0,288943658 | 0,844 | 0,645 | 1,36275E-91 |
| **UHRF1** | 5,74214E-55 | 0,289248631 | 0,544 | 0,328 | 1,4372E-50 |
| **LOXL1** | 4,60202E-77 | 0,289993185 | 0,825 | 0,652 | 1,15184E-72 |
| **CNN3** | 3,1738E-61 | 0,290926767 | 0,968 | 0,901 | 7,94371E-57 |
| **HIF1A** | 1,81243E-37 | 0,292122432 | 0,97 | 0,893 | 4,53632E-33 |
| **RPL7A** | 2,38416E-80 | 0,293869927 | 0,999 | 0,996 | 5,96731E-76 |
| **CENPE** | 9,58758E-22 | 0,294660773 | 0,53 | 0,39 | 2,39968E-17 |
| **CDH4** | 1,0922E-136 | 0,295928614 | 0,776 | 0,466 | 2,7336E-132 |
| **UBE2C** | 3,76312E-14 | 0,296011782 | 0,396 | 0,286 | 9,41871E-10 |
| **HMMR** | 3,61843E-16 | 0,296646503 | 0,363 | 0,254 | 9,05658E-12 |
| **TRAM2** | 1,58349E-57 | 0,298778275 | 0,879 | 0,687 | 3,96331E-53 |
| **RPS6** | 5,95804E-69 | 0,299275448 | 0,999 | 0,998 | 1,49124E-64 |
| **MXRA8** | 6,73493E-87 | 0,299837413 | 0,898 | 0,7 | 1,68569E-82 |
| **MYO1C** | 5,65565E-69 | 0,300017143 | 0,919 | 0,768 | 1,41555E-64 |
| **DNAJB4** | 8,59659E-50 | 0,301904364 | 0,936 | 0,828 | 2,15164E-45 |
| **SGO2** | 4,84031E-28 | 0,302030508 | 0,588 | 0,445 | 1,21148E-23 |
| **LMNA** | 8,20216E-43 | 0,302208489 | 0,976 | 0,932 | 2,05292E-38 |
| **PTGER2** | 7,388E-167 | 0,302412909 | 0,733 | 0,306 | 1,8491E-162 |
| **ANKRD28** | 2,98256E-63 | 0,302565427 | 0,903 | 0,715 | 7,46504E-59 |
| **DEK** | 1,23244E-10 | 0,304072561 | 0,954 | 0,877 | 3,08469E-06 |
| **MRC2** | 6,10323E-71 | 0,304220301 | 0,921 | 0,773 | 1,52758E-66 |
| **MIS18BP1** | 1,3814E-56 | 0,304251607 | 0,764 | 0,564 | 3,45751E-52 |
| **SULF1** | 2,7258E-112 | 0,305438385 | 0,627 | 0,261 | 6,8223E-108 |
| **C12orf57** | 1,4024E-91 | 0,305576535 | 0,974 | 0,87 | 3,51006E-87 |
| **HNRNPD** | 4,98535E-34 | 0,306234721 | 0,938 | 0,832 | 1,24778E-29 |
| **IGF2BP2** | 4,93545E-60 | 0,306899385 | 0,914 | 0,737 | 1,23529E-55 |
| **RPL10A** | 7,37882E-55 | 0,307774696 | 0,998 | 0,984 | 1,84684E-50 |
| **FBN1** | 9,0381E-46 | 0,307814529 | 0,91 | 0,764 | 2,26215E-41 |
| **MCM3** | 1,04064E-35 | 0,308155877 | 0,715 | 0,561 | 2,60463E-31 |
| **NAP1L1** | 2,0537E-26 | 0,308294605 | 0,991 | 0,983 | 5,1402E-22 |
| **CENPW** | 1,19538E-35 | 0,308963579 | 0,824 | 0,671 | 2,99193E-31 |
| **IQGAP1** | 8,09694E-61 | 0,312503228 | 0,945 | 0,848 | 2,02658E-56 |
| **DNMT1** | 2,00312E-38 | 0,313762051 | 0,896 | 0,74 | 5,01362E-34 |
| **CCNA2** | 7,51056E-47 | 0,313857223 | 0,471 | 0,252 | 1,87982E-42 |
| **ANXA2** | 1,76319E-36 | 0,314580728 | 1 | 0,996 | 4,41308E-32 |
| **ZBTB16** | 9,83426E-63 | 0,315406297 | 0,648 | 0,411 | 2,46142E-58 |
| **KIF20B** | 1,26473E-29 | 0,316767339 | 0,645 | 0,491 | 3,16549E-25 |
| **PLIN3** | 7,79064E-57 | 0,318651754 | 0,969 | 0,864 | 1,94992E-52 |
| **DTYMK** | 1,0891E-49 | 0,318696428 | 0,91 | 0,777 | 2,72592E-45 |
| **AC245297.3** | 3,45272E-74 | 0,319238529 | 0,91 | 0,779 | 8,64182E-70 |
| **MLPH** | 1,68777E-80 | 0,319765217 | 0,808 | 0,607 | 4,22432E-76 |
| **MME** | 4,29233E-74 | 0,321304126 | 0,923 | 0,759 | 1,07433E-69 |
| **USP1** | 4,26946E-45 | 0,323944246 | 0,798 | 0,63 | 1,0686E-40 |
| **CAVIN3** | 1,25396E-82 | 0,324257108 | 0,922 | 0,782 | 3,13854E-78 |
| **RRM1** | 4,83287E-41 | 0,324775434 | 0,846 | 0,686 | 1,20962E-36 |
| **HNRNPA3** | 1,34069E-50 | 0,327523533 | 0,977 | 0,941 | 3,35562E-46 |
| **CKAP4** | 3,75883E-52 | 0,327590842 | 0,95 | 0,867 | 9,40797E-48 |
| **XRCC5** | 1,02951E-56 | 0,327868123 | 0,978 | 0,942 | 2,57677E-52 |
| **HELLS** | 2,88847E-37 | 0,327933626 | 0,62 | 0,442 | 7,22954E-33 |
| **THY1** | 2,54069E-57 | 0,328552326 | 0,931 | 0,787 | 6,35909E-53 |
| **MYF6** | 3,2621E-113 | 0,328802515 | 0,77 | 0,378 | 8,1648E-109 |
| **CALM2** | 1,14261E-35 | 0,329762545 | 0,983 | 0,96 | 2,85984E-31 |
| **MT1E** | 1,8139E-19 | 0,330318066 | 0,988 | 0,928 | 4,54E-15 |
| **TLN1** | 2,77817E-69 | 0,330639788 | 0,968 | 0,902 | 6,95348E-65 |
| **VCAN** | 4,66157E-58 | 0,331913965 | 0,864 | 0,679 | 1,16674E-53 |
| **PLOD1** | 6,42199E-81 | 0,333505805 | 0,934 | 0,811 | 1,60736E-76 |
| **SOX4** | 9,10393E-74 | 0,336016333 | 0,783 | 0,564 | 2,27862E-69 |
| **CDKN3** | 1,38027E-21 | 0,340490824 | 0,461 | 0,344 | 3,45468E-17 |
| **TACC1** | 3,05458E-80 | 0,341332682 | 0,905 | 0,765 | 7,64532E-76 |
| **MT-ND3** | 2,99297E-42 | 0,342907607 | 0,987 | 0,947 | 7,49109E-38 |
| **WASF2** | 1,25683E-90 | 0,343496386 | 0,936 | 0,814 | 3,14571E-86 |
| **SLC38A1** | 1,15878E-69 | 0,344523018 | 0,891 | 0,702 | 2,9003E-65 |
| **GALNT1** | 2,7709E-104 | 0,345113009 | 0,913 | 0,735 | 6,9354E-100 |
| **EZR** | 1,12702E-75 | 0,34586031 | 0,917 | 0,756 | 2,82083E-71 |
| **ANLN** | 6,01604E-29 | 0,350724402 | 0,458 | 0,297 | 1,50575E-24 |
| **DLGAP5** | 5,91569E-26 | 0,35201518 | 0,36 | 0,213 | 1,48064E-21 |
| **TTC3** | 1,31679E-65 | 0,352823873 | 0,957 | 0,902 | 3,29579E-61 |
| **HNRNPU** | 1,05762E-47 | 0,353461354 | 0,958 | 0,858 | 2,64711E-43 |
| **GAS1** | 2,8574E-126 | 0,35391457 | 0,656 | 0,27 | 7,1517E-122 |
| **RHOBTB3** | 2,45015E-92 | 0,355723122 | 0,918 | 0,8 | 6,13247E-88 |
| **CCBE1** | 2,30585E-57 | 0,355776046 | 0,88 | 0,706 | 5,7713E-53 |
| **NNMT** | 9,82316E-68 | 0,356815382 | 0,98 | 0,914 | 2,45864E-63 |
| **RAD21** | 1,2595E-49 | 0,358868353 | 0,929 | 0,773 | 3,1524E-45 |
| **NASP** | 1,61635E-23 | 0,359618143 | 0,889 | 0,766 | 4,04557E-19 |
| **RBPJ** | 1,58141E-97 | 0,360159405 | 0,929 | 0,777 | 3,9581E-93 |
| **VKORC1** | 3,09923E-96 | 0,360899768 | 0,978 | 0,918 | 7,75706E-92 |
| **SMC2** | 1,0797E-40 | 0,361717476 | 0,872 | 0,709 | 2,70237E-36 |
| **CRIP2** | 4,9913E-110 | 0,361948234 | 0,894 | 0,681 | 1,2493E-105 |
| **PDGFA** | 3,2803E-120 | 0,36294777 | 0,826 | 0,475 | 8,2102E-116 |
| **TMPO** | 3,1398E-30 | 0,363079672 | 0,7 | 0,575 | 7,85859E-26 |
| **ARHGAP29** | 2,43527E-74 | 0,364043592 | 0,831 | 0,619 | 6,09523E-70 |
| **ZMYND8** | 1,60561E-86 | 0,365315267 | 0,88 | 0,672 | 4,01868E-82 |
| **MYOD1** | 1,5063E-118 | 0,365642277 | 0,551 | 0,185 | 3,77E-114 |
| **TRIM55** | 5,69311E-78 | 0,367135579 | 0,892 | 0,569 | 1,42493E-73 |
| **C1GALT1** | 1,19535E-81 | 0,368410798 | 0,895 | 0,735 | 2,99183E-77 |
| **MACF1** | 6,86338E-61 | 0,369233734 | 0,923 | 0,806 | 1,71784E-56 |
| **ENAH** | 9,67128E-89 | 0,369581246 | 0,932 | 0,794 | 2,42062E-84 |
| **KHDRBS1** | 1,85074E-81 | 0,370941015 | 0,934 | 0,819 | 4,63221E-77 |
| **SPATS2L** | 3,6504E-102 | 0,374549286 | 0,968 | 0,885 | 9,13648E-98 |
| **ASS1** | 2,18025E-66 | 0,374736983 | 0,841 | 0,596 | 5,45695E-62 |
| **AR** | 1,093E-167 | 0,374798213 | 0,83 | 0,52 | 2,7357E-163 |
| **MT-ND6** | 1,17566E-96 | 0,375650887 | 0,932 | 0,739 | 2,94255E-92 |
| **MT-ATP6** | 2,88559E-52 | 0,37663109 | 0,987 | 0,953 | 7,22235E-48 |
| **CBX1** | 1,3803E-72 | 0,377148552 | 0,926 | 0,805 | 3,45476E-68 |
| **CCDC88A** | 1,32736E-81 | 0,377699206 | 0,908 | 0,726 | 3,32225E-77 |
| **CAPG** | 2,68591E-47 | 0,378292055 | 0,654 | 0,512 | 6,72258E-43 |
| **PTTG1** | 5,09812E-17 | 0,380847987 | 0,809 | 0,7 | 1,27601E-12 |
| **CAV1** | 1,78809E-51 | 0,381226497 | 0,996 | 0,991 | 4,47542E-47 |
| **FLNA** | 2,90621E-76 | 0,384970071 | 0,96 | 0,871 | 7,27395E-72 |
| **CCNB1** | 7,82378E-15 | 0,387232168 | 0,622 | 0,519 | 1,95821E-10 |
| **LRP1** | 4,50419E-87 | 0,390321934 | 0,918 | 0,751 | 1,12735E-82 |

| | | | | | |
|---|---|---|---|---|---|
| EMP1 | 5,97066E-58 | 0,398240083 | 0,916 | 0,727 | 1,4944E-53 |
| TUBB3 | 4,72589E-63 | 0,398486895 | 0,895 | 0,76 | 1,18284E-58 |
| PBK | 1,99394E-34 | 0,398687247 | 0,402 | 0,23 | 4,99063E-30 |
| ALCAM | 2,35187E-43 | 0,401601297 | 0,927 | 0,773 | 5,8865E-39 |
| LTBP2 | 2,71405E-98 | 0,404131015 | 0,882 | 0,676 | 6,793E-94 |
| NFIC | 1,2894E-110 | 0,4061389 | 0,949 | 0,828 | 3,2271E-106 |
| RRM2 | 1,71002E-38 | 0,410804876 | 0,469 | 0,275 | 4,28002E-34 |
| PARP1 | 1,22844E-72 | 0,411401925 | 0,903 | 0,732 | 3,07467E-68 |
| HMGB3 | 1,0221E-33 | 0,414949219 | 0,83 | 0,716 | 2,55821E-29 |
| MGP | 2,52384E-42 | 0,417912596 | 0,148 | 0,019 | 6,31691E-38 |
| NUCKS1 | 7,50922E-51 | 0,418951767 | 0,983 | 0,941 | 1,87948E-46 |
| HMGN1 | 1,77486E-87 | 0,420421045 | 0,983 | 0,923 | 4,44231E-83 |
| TYMS | 2,19691E-36 | 0,420448284 | 0,733 | 0,592 | 5,49863E-32 |
| SPON2 | 1,5844E-155 | 0,421728179 | 0,677 | 0,266 | 3,9657E-151 |
| CXXC5 | 5,9064E-108 | 0,422530626 | 0,874 | 0,63 | 1,4783E-103 |
| TGFBI | 3,37865E-40 | 0,4230341 | 0,988 | 0,978 | 8,45643E-36 |
| P4HA1 | 8,39711E-93 | 0,424409551 | 0,947 | 0,853 | 2,10171E-88 |
| ENPP1 | 1,39311E-95 | 0,428172673 | 0,811 | 0,554 | 3,4868E-91 |
| HNRNPR | 6,72647E-73 | 0,430057025 | 0,943 | 0,816 | 1,68357E-68 |
| SMC4 | 1,75104E-32 | 0,432869148 | 0,831 | 0,678 | 4,38268E-28 |
| GABPB1-AS1 | 2,3763E-130 | 0,434200398 | 0,867 | 0,619 | 5,9475E-126 |
| COL5A2 | 4,82394E-80 | 0,434827461 | 0,909 | 0,734 | 1,20738E-75 |
| CKS1B | 6,68406E-36 | 0,43538577 | 0,896 | 0,78 | 1,67295E-31 |
| HIST1H1B | 1,69937E-28 | 0,435567437 | 0,268 | 0,123 | 4,25335E-24 |
| SLC20A1 | 3,91583E-61 | 0,435950587 | 0,9 | 0,742 | 9,80094E-57 |
| NEAT1 | 1,47786E-53 | 0,436359127 | 0,937 | 0,798 | 3,69893E-49 |
| EPB41L2 | 2,76089E-84 | 0,437229139 | 0,903 | 0,779 | 6,91024E-80 |
| PTMS | 1,2567E-114 | 0,437659125 | 0,991 | 0,963 | 3,1454E-110 |
| RPL3 | 5,31246E-97 | 0,438843413 | 0,999 | 0,991 | 1,32966E-92 |
| VIM | 9,87601E-55 | 0,439894559 | 1 | 0,996 | 2,47187E-50 |
| CLSPN | 5,69398E-31 | 0,447186971 | 0,495 | 0,335 | 1,42515E-26 |
| CRLF1 | 1,5679E-175 | 0,448566729 | 0,77 | 0,317 | 3,9243E-171 |
| FBLN1 | 1,8176E-212 | 0,450068817 | 0,776 | 0,305 | 4,5492E-208 |
| CARHSP1 | 4,80902E-61 | 0,453071103 | 0,914 | 0,785 | 1,20365E-56 |
| MAT2A | 1,5897E-105 | 0,459190075 | 0,925 | 0,773 | 3,9787E-101 |
| DDIT4 | 4,86878E-83 | 0,46360492 | 0,77 | 0,498 | 1,21861E-78 |
| PRC1 | 1,14537E-30 | 0,466173793 | 0,564 | 0,408 | 2,86674E-26 |
| TGM2 | 3,7776E-09 | 0,466562077 | 0,739 | 0,666 | 9,45495E-05 |
| CELF2 | 3,3427E-111 | 0,471207527 | 0,822 | 0,539 | 8,3665E-107 |
| OLFML3 | 5,77766E-74 | 0,471303976 | 0,77 | 0,53 | 1,44609E-69 |
| ZFP36L1 | 5,22389E-64 | 0,472109113 | 0,922 | 0,762 | 1,30749E-59 |
| CKAP2 | 5,32186E-49 | 0,47461902 | 0,788 | 0,615 | 1,33201E-44 |
| RRBP1 | 8,39027E-89 | 0,474676256 | 0,949 | 0,821 | 2,1E-84 |
| HEY1 | 7,309E-148 | 0,476247508 | 0,695 | 0,331 | 1,8294E-143 |
| ASPM | 5,58142E-29 | 0,482848502 | 0,38 | 0,218 | 1,39697E-24 |
| EPS8 | 9,8573E-115 | 0,484787624 | 0,944 | 0,815 | 2,4672E-110 |
| CDK2AP1 | 7,929E-156 | 0,486561809 | 0,973 | 0,9 | 1,9845E-151 |
| STK17B | 8,3292E-131 | 0,488131687 | 0,885 | 0,672 | 2,0847E-126 |
| SLC38A2 | 1,39595E-90 | 0,496507883 | 0,922 | 0,757 | 3,49391E-86 |
| CDC42EP3 | 1,00061E-92 | 0,50194127 | 0,909 | 0,732 | 2,50442E-88 |
| TSC22D3 | 1,30504E-93 | 0,509121205 | 0,921 | 0,723 | 3,26638E-89 |
| NUSAP1 | 2,5362E-33 | 0,512178985 | 0,475 | 0,313 | 6,34785E-29 |
| LMO7 | 2,35293E-97 | 0,514817256 | 0,927 | 0,77 | 5,88915E-93 |
| CBX5 | 5,70976E-60 | 0,518527876 | 0,918 | 0,797 | 1,4291E-55 |
| SNAI2 | 8,0268E-124 | 0,519682387 | 0,898 | 0,679 | 2,009E-119 |
| TPX2 | 5,57323E-46 | 0,527637239 | 0,672 | 0,468 | 1,39492E-41 |
| PGF | 1,2431E-112 | 0,533663692 | 0,807 | 0,492 | 3,1114E-108 |
| STMN1 | 4,26523E-08 | 0,543120474 | 0,939 | 0,925 | 0,001067544 |
| MSI2 | 1,1666E-128 | 0,543310459 | 0,902 | 0,736 | 2,9199E-124 |
| BIRC5 | 4,46197E-33 | 0,552686153 | 0,489 | 0,329 | 1,11679E-28 |
| LSP1 | 4,2549E-145 | 0,55717755 | 0,939 | 0,658 | 1,065E-140 |
| TUBB | 4,5321E-63 | 0,565553137 | 0,973 | 0,916 | 1,13434E-58 |
| HEYL | 3,0597E-163 | 0,566494603 | 0,734 | 0,327 | 7,6582E-159 |
| TUBA1B | 1,58651E-45 | 0,571808776 | 0,977 | 0,922 | 3,97088E-41 |
| HMGB2 | 2,34179E-26 | 0,576568438 | 0,797 | 0,665 | 5,86125E-22 |
| AHNAK | 9,49921E-95 | 0,587489931 | 0,971 | 0,889 | 2,37756E-90 |
| HMGB1 | 7,91704E-31 | 0,599574285 | 0,991 | 0,974 | 1,98155E-26 |
| MKI67 | 1,19162E-30 | 0,602733065 | 0,395 | 0,229 | 2,98251E-26 |
| LOX | 8,92482E-40 | 0,604512141 | 0,872 | 0,739 | 2,23379E-35 |
| UGP2 | 8,985E-57 | 0,607602481 | 0,968 | 0,897 | 2,24886E-52 |
| MT1M | 1,79589E-52 | 0,617813668 | 0,679 | 0,435 | 4,49494E-48 |
| NR2F1 | 3,1923E-135 | 0,634127749 | 0,793 | 0,506 | 7,9899E-131 |
| MARCKS | 2,057E-128 | 0,647932384 | 0,931 | 0,811 | 5,1484E-124 |
| PRKDC | 1,7075E-124 | 0,653667393 | 0,933 | 0,831 | 4,2737E-120 |
| PTMA | 5,98777E-75 | 0,655726473 | 0,991 | 0,963 | 1,49868E-70 |
| SIX1 | 1,828E-182 | 0,662590597 | 0,879 | 0,627 | 4,5753E-178 |
| PCLAF | 3,71391E-42 | 0,691057925 | 0,569 | 0,381 | 9,29555E-38 |
| CRYAB | 3,28663E-09 | 0,696726543 | 0,972 | 0,902 | 8,2261E-05 |
| FKBP5 | 5,3591E-213 | 0,711079502 | 0,945 | 0,754 | 1,3413E-208 |
| DKK1 | 6,34517E-57 | 0,752331451 | 0,916 | 0,737 | 1,58813E-52 |
| COL1A1 | 5,71371E-71 | 0,761229279 | 0,981 | 0,925 | 1,43009E-66 |
| PCDH9 | 1,2218E-120 | 0,786040163 | 0,913 | 0,707 | 3,0581E-116 |
| TOP2A | 3,83947E-55 | 0,795153013 | 0,537 | 0,292 | 9,60981E-51 |
| COL3A1 | 1,1594E-102 | 0,80349378 | 0,892 | 0,716 | 2,90191E-98 |
| FN1 | 4,22394E-92 | 0,807870002 | 0,991 | 0,958 | 1,05721E-87 |
| NPTX2 | 1,1982E-152 | 0,814974946 | 0,872 | 0,539 | 2,9989E-148 |
| COL1A2 | 1,0966E-108 | 0,892639088 | 0,973 | 0,864 | 2,7447E-104 |
| HES1 | 2,7208E-151 | 0,920114569 | 0,92 | 0,741 | 6,8099E-147 |
| IFITM2 | 4,6778E-237 | 0,930181869 | 0,991 | 0,888 | 1,1708E-232 |
| TUBA1A | 1,0638E-125 | 0,959611761 | 0,952 | 0,829 | 2,6627E-121 |
| ID3 | 1,4939E-155 | 0,968175924 | 0,944 | 0,772 | 3,7392E-151 |
| MYF5 | 1,9548E-149 | 1,02515735 | 0,827 | 0,464 | 4,8926E-145 |
| ATP2B1 | 5,7273E-141 | 1,025835673 | 0,954 | 0,858 | 1,4335E-136 |
| CENPF | 2,86662E-46 | 1,060132233 | 0,583 | 0,381 | 7,17487E-42 |
| SERPINE2 | 8,04069E-66 | 1,069346241 | 0,954 | 0,832 | 2,0125E-61 |
| HIST1H4C | 3,34577E-69 | 1,260371915 | 0,896 | 0,706 | 8,37412E-65 |
| ID1 | 5,9318E-246 | 1,409596442 | 0,926 | 0,701 | 1,4847E-241 |

**Supplemental table 2.** List of differentially expressed genes (DEG) between control and DM1 patients myoblasts showing the p-value (p_val), the average log2 fold-change (avg_log2FC), the proportion of cells expressing each gene in control and patients, and the adjusted p-value (adj_p_val).

| Characteristics | Total (n=103) | Low IL-6 level (n=80) | High IL-6 level (n=23) |
|---|---|---|---|
| **Age, y** | | | |
| Mean (SD) | 43.3 (10.1) | 43.0 (10.7) | 43.2 (7.8) |
| [min-max] | [20-77] | [20-77] | [30-69] |
| **Sex, n (%)** | | | |
| Men | 38 (36.9) | 29 (36.3) | 9 (39.1) |
| Women | 65 (63.1) | 51 (63.7) | 14 (60.9) |
| **Body mass index** | | | |
| Mean (SD) | 25.4 (5.5) | 24.1 (4.6) | 28.7 (6.7) |
| [min-max] | [14.5-43.0] | [14.5-41.4] | [16.4-43.0] |
| **Phenotype, n (%)** | | | |
| Late-onset | 20 (19.4) | 17 (21.3) | 3 (13.0) |
| Adult | 83 (80.6) | 63 (78.8) | 20 (87.0) |
| **CTG repeat length** | | | |
| Mean (SD) | 576.1 (378.1) | 575.3 (399.5) | 578.8 (299.8) |
| [min-max] | [59-2000] | [59-2000] | [63-1089] |
| **Disease duration (n = 74), y** | | | |
| Mean (SD) | 19.7 (7.9) | 19.5 (8.0) | 20.3 (7.96) |
| [min-max] | [3-35] | [3-35] | [6-34] |
| **TNF-α level** | | | |
| Mean (SD) | 1.67 (1.61) | 1.68 (1.81) | 1.63 (0.43) |
| [min-max] | [0.47-16.06] | [0.47-16.06] | [1.00-2.42] |
| **IL-6 level** | | | |
| Mean (SD) | 3.28 (3.25) | 2.04 (0.84) | 7.61 (4.61) |
| [min-max] | [0.72-23.49] | [0.72-4.11] | [4.48-23.49] |

**Supplemental table 3.** Clinical characteristics of DM1 patients that were subjected to functional capacity tests and assessment of serum IL-6 levels.

| Test | R2, F value (df; Residual), p-value | Variables (Standardized B, p-value) |
|---|---|---|
| Timed-Up and Go | 0.170, F (3, 99) = 6.77, p <.001 | Phenotype (0.39, p<.001)<br>Age (0.35, p=.002)<br>**IL-6 (0.18, p=.048)** |
| 10mWT | 0.299, F (4, 98) = 10.46, p<.001 | Phenotype (-0.48, p<.001)<br>Sex (-0.27, p=.002)<br>**IL-6 (-0.2, p=.003)**<br>Age (-0.23, p=.026) |
| Berg Balance Scale | 0.133, F (2, 70) = 5.37, p=.007 | **IL-6 (-0.30, p=.010)**<br>Phenotype (-0.22, p=.050) |
| Pinch test | 0.492, F (2, 100) = 48.51, p<.001) | Phenotype (-0.64, p<.001)<br>Sex (-0.29, p<.001) |
| Ankle dorsiflexors | 0.349, F (3, 98) = 17.49, p<.001) | Phenotype (-0.42, p<.001)<br>Sex (0.31, p<.001)<br>**IL-6 (-0.24, p=.005)** |
| Hip flexors | 0.305, F (3, 98) = 14.34, p<.001) | Phenotype (-0.55, p<.001)<br>**IL-6 (-0.27, p=.002)**<br>Age (-0.32, p=.002) |
| Knee extensors | 0.138, F (2, 99) = 7.95, p<.001) | Phenotype (-0.29, p=.003)<br>**IL-6 (-0.20, p=.034)** |
| Knee flexors | 0.197, F (2, 100) = 12.23, p<.001) | Phenotype (-0.38, p<.001)<br>**IL-6 (-0.19, p=.034)** |
| Shoulder flexors | 0.244, F (2, 97) = 15.65, p<.001) | Phenotype (-0.39, p<.001)<br>**IL-6 (-0.28, p=.002)** |
| Shoulder abductors | 0.327, F (2, 98) = 23.76, p<.001) | Phenotype (-0.51, p<.001)<br>**IL-6 (-0.22, p=.009)** |
| Elbow flexors | 0.347, F (2, 99) = 26.29, p<.001) | Phenotype (-0.55, p<.001)<br>**IL-6 (-0.18, p=.034)** |
| Elbow extensors | 0.441, F (3, 98) = 19.09, p<.001) | Phenotype (-0.56, p<.001)<br>Sex (0.31, p<.001)<br>Age (-0.32, p<.001)<br>**IL-6 (-0.206, p<.001)** |

**Supplemental table 4.** Step wise regression model of factors predicting muscle strength and functional outcomes in DM1 patients.

| Gene | Forward sequence | Reverse sequence |
|---|---|---|
| *P16* | TGTTCGCATTGCCAAGGTTC | CGTTTCCGTGAATGTTGTCCC |
| *P21* | CTGGGGATGTCCGTCAGAAC | GTGACAGGTCCACATGGTCT |
| *RPLP0* | GGCAGCATCTACAACCCTGA | CAGGACTCGTTTGTACCCGT |

**Supplemental table 5.** List of primers used for qPCR experiments
